# Prenatal exposure to dietary levels of glyphosate disrupts metabolic, immune, and behavioral markers across generations in mice

**DOI:** 10.1101/2024.08.27.609990

**Authors:** J.A. Barnett, J.K. Josephson, N. Haskey, M.M. Hart, K.K. Soma, A. Verdugo, C.J. McComb, M.L. Bandy, S. Ghosh, C. Letef, A. Copp, R. Ishida, E. Yuzbashian, J. Gibon, J. Ye, R.T. Giebelhaus, S.J. Murch, M.M. Jung, D.L. Gibson

**Affiliations:** Department of Biology, University of British Columbia; Kelowna, V1V 1V7, Canada; Department of Zoology, University of British Columbia; Vancouver, V6T 1Z4, Canada; Djavad Mowafaghian Centre for Brain Health, University of British Columbia; Vancouver, V6T 1Z3, Canada; Department of Psychology, University of British Columbia; Vancouver, V6T 1Z4, Canada; Department of Chemistry, University of British Columbia; Kelowna, V1V 1V7, Canada; Department of Chemistry, University of Alberta; Edmonton, T6G 2N4, Canada; Faculty of Medicine, University of British Columbia; Kelowna, V1V 1V7, Canada

## Abstract

Glyphosate, a widely used herbicide in North America, has become prevalent in the food supply, raising concerns about potential health impacts. In this exploratory study, male and female F0 mice were exposed to glyphosate through drinking water during mating and gestation. We investigated whether prenatal exposure at dietary-relevant levels (0.01 mg/kg/day, Average American Diet, [AAD]) or the U.S. EPA’s acceptable daily intake (1.75 mg/kg/day, [EPA upper limit]) altered gut, metabolic, and behavioral outcomes across two generations in mice with or without genetic susceptibility to colitis (*Muc2*^+/-^ and *Muc2*^-/-^, respectively). Healthy (*Muc2*^+/-^) offspring of glyphosate-exposed mice exhibited colonic goblet cell depletion, reduced mucin-2 expression, and pro-inflammatory cytokine profiles in both F1 and F2 generations. These healthy (*Muc2*^+/-^) offspring also developed metabolic dysfunction, including impaired glucose tolerance, insulin resistance, and reduced GLP-1 in serum. Behavioral deficits were also observed in healthy (*Muc*^+/-^) mice including reduced locomotion and working memory, and these changes were associated with altered microbiome composition and gut-brain mediators. These findings suggest that prenatal glyphosate exposure, even below regulatory thresholds, may disrupt multiple physiological systems across generations, highlighting the need for further research and regulatory consideration.

## Introduction

Glyphosate (Roundup®) is one of North America’s most widely used herbicides, with over 160 million kilograms applied annually.^1,2^ Agricultural practices, including the development of glyphosate-resistant crops and glyphosate’s use as a pre-harvest desiccant, have led to increased residues in commonly consumed foods such as wheat, corn, soy, and oats. ³ These ingredients form the basis of the Western diet, which is frequently linked to chronic inflammatory diseases.^4-6^ While poor nutritional quality is often blamed, less attention has been given to the environmental contaminants embedded within this diet—particularly pesticide residues like glyphosate.

Glyphosate targets the Shikimate pathway, which is found in plants, fungi, and bacteria but is absent in mammals.^7^ This absence led to the assumption that glyphosate was biologically inert in humans. However, emerging evidence suggests that glyphosate may indirectly affect host physiology by altering the gut microbiota,^7,8^ modulating immune responses,^9^ and disrupting endocrine signaling.^10^ These concerns are amplified during prenatal and early-life exposures when developmental systems are particularly sensitive to environmental cues. In addition to vertical transmission of microbes from mother to infant through delivery and breastfeeding, changes in maternal immunity, hormonal signaling, and microbiota-derived metabolites can shape fetal development through placental and lactational transfer, increasing the long-term risk for metabolic,^11^ immune,^12^ and neurodevelopmental disorders.^13,14^

Most studies investigating glyphosate’s biological effects rely on doses far exceeding those encountered through diet, limiting their real-world applicability.^7,8^ Additionally, prior work has largely focused on direct exposure in healthy subjects, with little attention given to vulnerable populations or the possibility of heritable effects. It remains unclear whether glyphosate levels comparable to those found in the food supply can impact offspring health— particularly in the context of genetic susceptibility to disease. To address this gap, we examined whether prenatal glyphosate exposure at human-relevant levels—0.01 mg/kg/day, reflecting estimated dietary intake (Average American Diet, AAD), and 1.75 mg/kg/day, the U.S. EPA’s acceptable daily intake (EPA Upper Limit)—disrupts immune, metabolic, and neurobehavioral outcomes across two generations in both healthy (*Muc2*^+/-^) and colitis-susceptible (*Muc2*^-/-^) mice. Based on previous studies linking glyphosate to microbiome disruption, we hypothesized that prenatal exposure at these doses would alter microbial composition and function, leading to increased intestinal inflammation, metabolic dysfunction, and behavioral changes that persist across generations in both genotypes.

## Results

### Prenatal glyphosate exposure induces microscopic colitis in healthy (*Muc2*^+/-^) offspring and promotes intestinal inflammation

To assess how prenatal glyphosate exposure affects intestinal pathology, we examined markers of morbidity and inflammation in F1 and F2 offspring from healthy (*Muc2*^+/-^) and colitis-susceptible (*Muc2*^-/-^) mice. Although glyphosate-exposed healthy offspring did not exhibit overt colitis, histopathological analysis revealed morphological changes across both generations and exposure levels, consistent with microscopic colitis (Fig. 1A). Goblet cell depletion and crypt hyperplasia were the primary drivers of pathology (Fig. 1B–C), with goblet cell loss accompanied by reduced mucin-2 expression in the colon (Fig. 1C), indicating compromised mucus barrier function often seen in IBD patients.^15^

**Figure 1.**
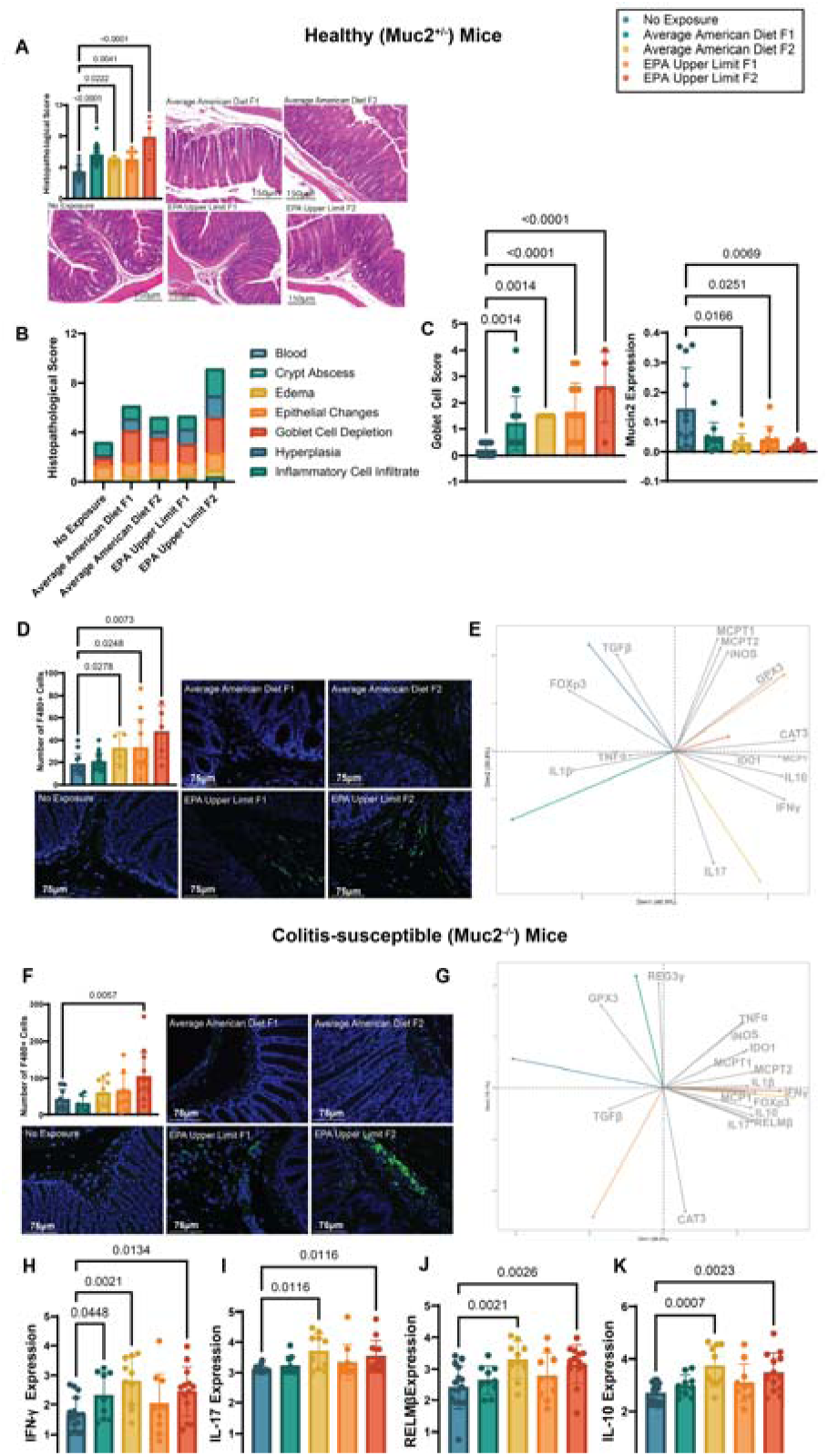
Prenatal glyphosate exposure induces microscopic colitis and colonic inflammation across generations in mice. **(A)** Histopathological scoring of colon tissue reveals increased damage in healthy (*Muc2*^+/-^) F1 and F2 offspring of glyphosate-exposed parents compared to non-exposed animals. Damage was scored across crypt architecture, goblet cell abundance, and immune cell infiltration (Kruskal-Wallis, *p* < 0.0001, η² = 0.493, FDR-corrected; n = 5–16 per group.) **(B–C)** Goblet cell depletion and epithelial hyperplasia were the primary contributors to histological scores (Two-way ANOVA, *p* = 0.0001, FDR-corrected.) These changes were associated with decreased mucin-2 expression in colon tissue (Kruskal-Wallis, *p* = 0.025, η² = 0.178.) **(D)** Increased macrophage (F4/80+) infiltration was observed in the colonic submucosa of EPA F2 healthy (*Muc2*^+/-^) offspring (Kruskal-Wallis, *p* = 0.015, η² = 0.135.) **(E)** Principal component analysis (PCA) of colonic cytokine expression shows a pro-inflammatory shift in F1 and F2 glyphosate-exposed offspring, including enrichment of TH1/TH17 cytokines (e.g., TNF-α, IL-17, IFN-γ.) **(F–K)** In colitis-susceptible (*Muc2*^-/-^) offspring, glyphosate exposure induced modest increases in colonic inflammation, including elevated macrophage infiltration and increased expression of IFN-γ, IL-17, IL-10, and RELMβ (ANOVA/Kruskal-Wallis, FDR-corrected.) **Group Definitions:** No exposure = no glyphosate exposure in F0; AAD = Average American Diet intake level (0.01mg/kg/day); EPA upper limit = U.S. EPA acceptable daily intake (1.75mg/kg/day.) “F1” and “F2” refer to first- and second-generation offspring of F0-exposed breeders. **Genotypes:** *Muc2*^+/-^ = healthy mice; *Muc2*^-/-^ = colitis-susceptible mice. **Sex:** Sex was not found to be a driving factor in histopathological damage (PERMANOVA, *p*=0.268), goblet-cell depletion (PERMANOVA, *p*=0.485) or macrophage infiltration (PERMANOVA, *p*=0.330) in healthy mice, however sex was found to influence mucin-2 gene expression (PERMANOVA, *p*=0.019. In colitis-susceptible mice, sex was not found to be a driving factor in macrophage infiltration (PERMANOVA, *p*=0.564), IFN-γ (PERMANOVA, *p*=0.604), RELMβ (PERMANOVA, *p*=0.395), IL-17 (PERMANOVA, *p*=0.490), or IL-10 (PERMANOVA, *p*=0.765) gene expression. Primer sequences used for cytokine analysis are found in table S5. **Statistical Summary:** Confidence intervals, and sample sizes broken down by sex provided in table S6.

EPA F2 healthy (*Muc2*^+/-^) offspring also showed increased F4/80+ macrophage infiltration in the colonic submucosa (Fig. 1D.) Cytokine expression patterns, analyzed via PCA, showed distinct inflammatory signatures by treatment group (Fig. 1E) F1 offspring of AAD-exposed parents were characterized by a TNF-α and IL-1β–driven response, while F2 offspring shifted toward a TH17-like cytokine profile with enrichment in IFN-γ and IL-17. EPA-exposed groups showed upregulation of oxidative stress markers (GPX2, CAT3) and mast cell proteases (MCPT1, MCPT2), consistent with innate immune activation.

In colitis-susceptible (*Muc2*^-/-^) offspring, prenatal glyphosate exposure had a more limited effect. EPA F2 mice showed elevated macrophage infiltration (Fig. 1F). PCA revealed a similar pro-inflammatory cytokine profile (Fig. 1G), including increased expression of IFN-γ, and IL-17 (Fig. 1 H–K), but changes were modest compared to healthy (*Muc2*^+/-^) littermates —potentially reflecting a ceiling effect due to baseline inflammation. Colitis-susceptible (*Muc2*^-/-^) offspring also showed increased expression of the goblet cell mediator RELM-β, which has been shown to be a driving force in the development of colitis within this model.^16^

These findings suggest that prenatal glyphosate exposure induces microscopic inflammation and mucus barrier dysfunction in healthy (*Muc2*^+/-^) offspring, with effects that persist across generations. In colitis-susceptible (*Muc2*^-/-^) mice, baseline inflammation may mask additional glyphosate-driven pathology.

### Prenatal Glyphosate Exposure Impairs Glucose Metabolism, Reduces Insulin Sensitivity, and Alters Metabolic Hormone Production in Healthy Mice, with Potential Links to Endotoxemia

Given the increased risk of metabolic syndrome in individuals with IBD,^17^ we next assessed whether prenatal glyphosate exposure disrupted metabolic regulation in healthy (*Muc2*^+/-^) and colitis-susceptible (*Muc2*^-/-^) offspring. In the oral glucose tolerance test (OGTT), EPA-exposed healthy (*Muc2*^+/-^) F2 offspring showed impaired glucose clearance, with elevated blood glucose 15 minutes post-gavage and increased AUC (Kruskal-Wallis, *p* = 0.0106, η² = 0.129; Fig. 2A). No such impairment was observed in the AAD group, suggesting a possible threshold effect for glyphosate-induced glycemic disruption in the F2 generation. In the insulin tolerance test (ITT), EPA-exposed F1 healthy (*Muc2*^+/-^) mice showed reduced insulin sensitivity relative to non-exposed mice (ANOVA, *p* = 0.0326, η²p = 0.174; Fig. 2B.) Serum GLP-1 levels were significantly reduced in both AAD- and EPA-exposed F1 and F2 offspring, with a trend toward reduction in AAD F2 mice (Kruskal-Wallis, *p* = 0.015, η² = 0.145; Fig. 2C.) Ghrelin levels were lower in F1 mice, while leptin was elevated in AAD F1/F2 and EPA F1 offspring (Fig. 2D–E), suggesting dysregulated appetite control and possible leptin resistance.^18,19^ This hormonal profile mirrors patterns observed in diet-induced obesity^20^ and gut permeability models.^21,22^

**Figure 2.**
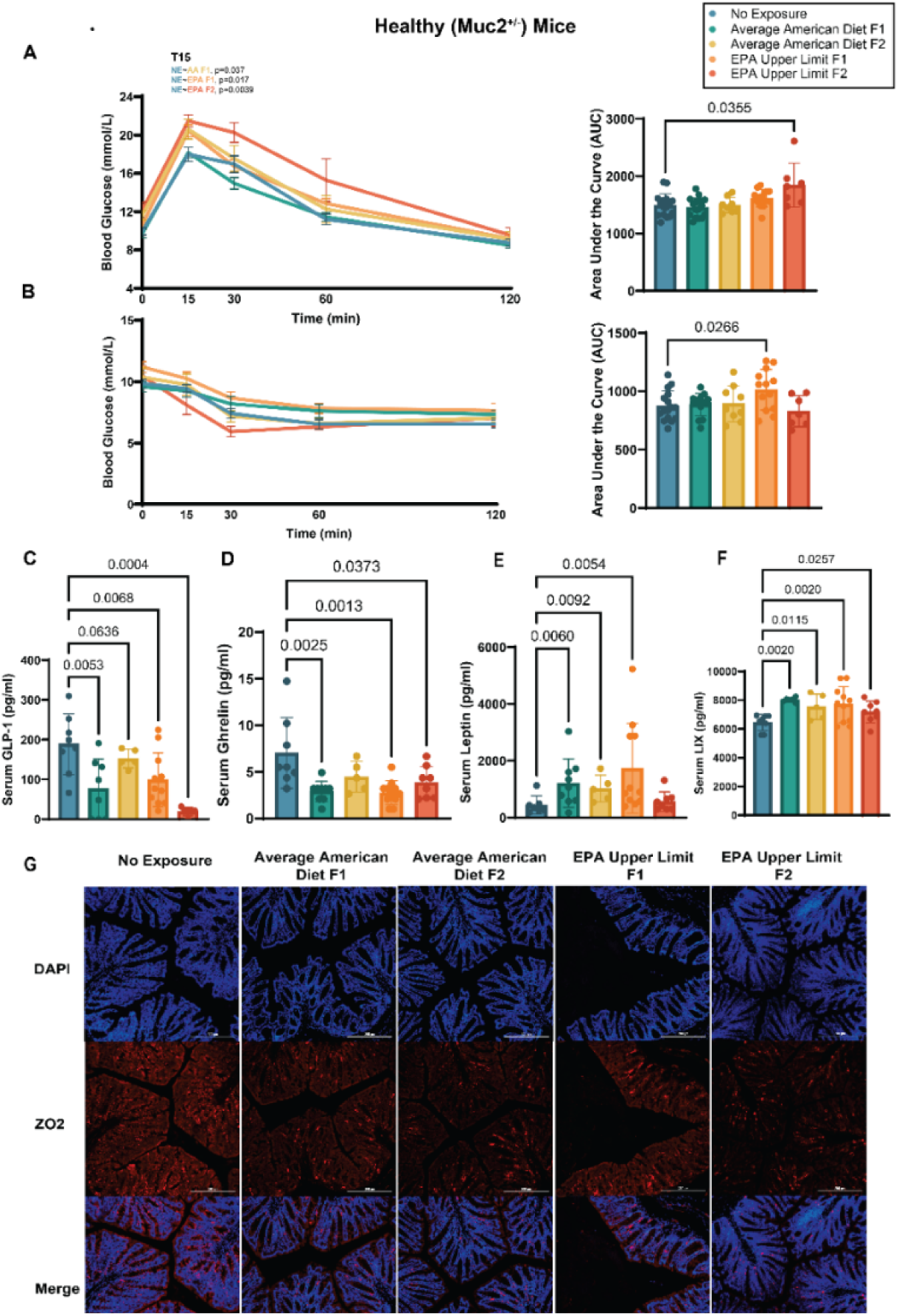
Prenatal glyphosate exposure impairs glucose regulation and alters metabolic hormones in healthy (*Muc2*^+/-^) offspring across generations. **(A)** Oral glucose tolerance test (OGTT) in F2 offspring reveals impaired glucose clearance in EPA-exposed healthy (*Muc2*^+/-^) mice compared to non-exposed controls (Kruskal-Wallis, *p* = 0.0106, η² = 0.129, FDR-corrected; n = 10–12 per group.) **(B)** Insulin tolerance test (ITT) in F1 healthy (*Muc2*^+/-^) offspring shows reduced insulin sensitivity in the EPA group (ANOVA, *p* = 0.0326, η²p = 0.174, FDR-corrected). **(C)** GLP-1 levels were significantly reduced in both AAD- and EPA-exposed F1 and F2 *Muc2*^+/-^ mice (Kruskal-Wallis, *p* = 0.015, η² = 0.145, FDR-corrected.) **(D)** Ghrelin levels were decreased in F1 healthy (*Muc2*^+/-^) offspring following prenatal glyphosate exposure. **(E)** Leptin levels were elevated in AAD F1/F2 and EPA F1 offspring, suggesting potential leptin resistance. **(F)** Serum levels of LIX (CXCL5), a neutrophil chemoattractant linked to intestinal permeability and inflammation, were elevated in glyphosate-exposed offspring. **(G)** Immunofluorescent staining for ZO-2 revealed disrupted tight junction organization in the colonic epithelium of EPA F2 mice, indicative of barrier dysfunction. **Group Definitions:** No exposure = no glyphosate exposure in F0; AAD = Average American Diet intake level (0.01 mg/kg/day); EPA Upper Limit = U.S. EPA acceptable daily intake (1.75 mg/kg/day.) “F1” and “F2” refer to first- and second-generation offspring of F0-exposed breeders. Genotypes: *Muc2*^+/-^ = healthy mice; *Muc2*^-/-^ = colitis-susceptible mice (not shown.) **Sex:** Sex was found to influence glucose tolerance (PERMANOVA, *p*=0.042), and may influence insulin tolerance (PERMANOVA, *p*=0.062). Sex was not found to be a driving influence in the observed changes in serum GLP-1 (PERMANOVA, *p*=0.368), ghrelin (PERMANOVA, *p*=0.791), leptin (PERMANOVA, *p*=0.072) or LIX (PERMANOVA, *p*=0.823). **Statistical Summary:** Confidence intervals, and sample sizes broken down by sex provided in table S6.

To explore whether these metabolic changes were linked to gut barrier dysfunction, we measured markers of endotoxemia and tight junction integrity. Serum LIX (CXCL5) levels were elevated in healthy (*Muc2*^+/-^) offspring of glyphosate-exposed mice (Fig. 2F), consistent with systemic immune activation. In the colon, disorganized ZO-2 protein expression in EPA F2 mice, suggesting tight junction disruption (Fig. 2G.) These findings indicate compromised barrier function as a potential upstream driver of metabolic dysregulation. Like what was observed in intestinal pathology, colitis-susceptible (*Muc2*^-/-^) offspring showed no glyphosate-related changes in glucose tolerance, insulin sensitivity, or hormone levels, again supporting a potential ceiling effect, where underlying inflammation and metabolic stress within the model eclipses additional glyphosate effects.

Together, these findings show that prenatal glyphosate exposure—at doses reflective of human dietary intake—disrupts metabolic regulation in healthy (*Muc2*^+/-^) mice across generations. These effects likely arise through gut-mediated mechanisms involving barrier dysfunction and immune activation and appear less pronounced in mice with a preexisting susceptibility to intestinal inflammation (*Muc2*^-/-^ mice.)

### Prenatal glyphosate exposure reduces locomotor activity, impairs working memory and alters the gut-brain axis in offspring

Individuals with IBD face an elevated risk of psychological and cognitive dysfunction.^23-25^ To investigate whether glyphosate contributes to these outcomes across generations, we assessed behavior and gut-brain signaling in offspring following prenatal exposure. Glyphosate-exposed healthy (*Muc2*^+/-^) offspring showed reduced locomotor activity, impaired working memory, and altered neuroimmune markers, particularly in the F2 generation. In contrast, colitis-susceptible (*Muc2*^-/-^) offspring showed no overt behavioral changes but did exhibit molecular signatures of enteric neuroinflammation.

In the open field test, F2 EPA-exposed *Muc2*^+/-^ mice travelled less distance and had lower average body velocity (*p* = 0.0003, η²p = 0.460; Fig. 3A), indicating reduced locomotor activity. These mice also showed fewer unsupported rearings (*p* = 0.018, η²p = 0.219; Fig. 3B), which may reflect diminished exploratory drive or subtle motor deficits. Anxiety-like behavior remained unchanged across exposure groups, based on center zone time, light/dark box preference, and serum corticosterone (fig. S3) Working memory was impaired in F1 AAD-exposed *Muc2*+/– mice, which made more errors in the radial arm maze (*p* = 0.041, η²p = 0.147; Fig. 3C) These deficits coincided with reduced serum kynurenine in both AAD F1 and EPA F2 offspring (*p* = 0.024, η² = 0.182; Fig. 3D) Given kynurenine’s role in generating neuroactive metabolites, this may signal disrupted neuroprotective metabolism.

**Figure 3.**
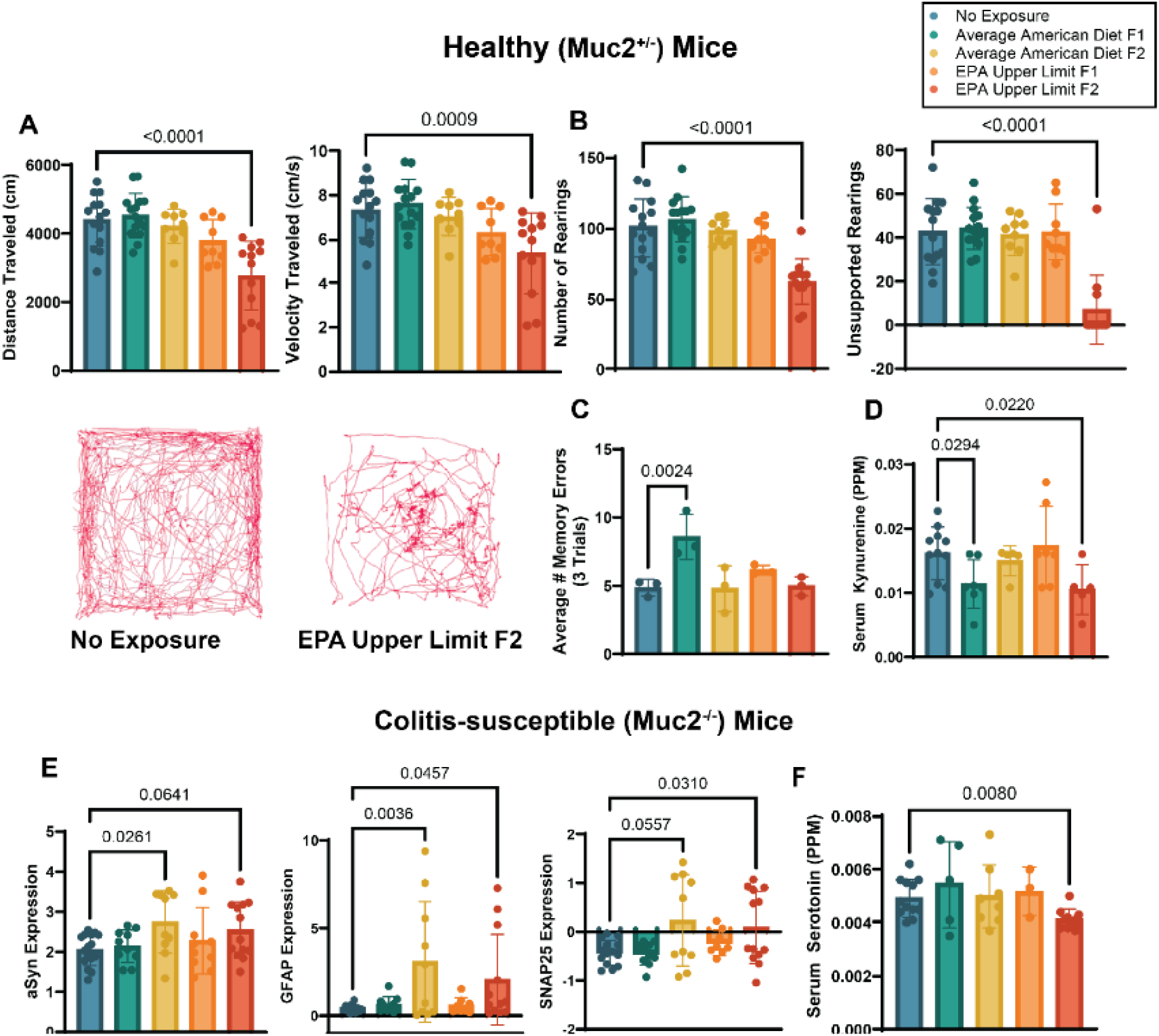
Prenatal glyphosate exposure alters locomotor activity, working memory, and gut–brain axis signaling in healthy (*Muc2*^+/-^) and colitis-susceptible (*Muc2*^-/-^) offspring. **(A)** Healthy (*Muc2*^+/-^) EPA F2 offspring exhibited significantly reduced distance traveled and body velocity in the open field test (*p* = 0.0003, η²p = 0.460, FDR-corrected; n = 6–10/group.) **(B)** Unsupported rearing frequency was significantly reduced in the same group (*p* = 0.018, η²p = 0.219), suggesting impaired exploratory behavior. **(C)** AAD F1 healthy (*Muc2*^+/-^) offspring displayed impaired working memory, reflected by increased radial arm maze errors (*p* = 0.041, η²p = 0.147). **(D)** Serum kynurenine levels were significantly reduced in AAD F1 and EPA F2 healthy (*Muc2*^+/-^) offspring (*p* = 0.024, η² = 0.182), consistent with disruption of neuroprotective metabolic pathways. **(E)** F2 colitis-susceptible (*Muc2*^-/-^) offspring exhibited elevated expression of α-synuclein, GFAP, and SNAP-25 in colonic tissue following AAD and EPA exposure (all *p* < 0.05, FDR-corrected), suggesting enteric neuroinflammation. **(F)** Serum serotonin was significantly reduced in EPA F2 *Muc2*^-/-^ offspring (*p* = 0.031, η² = 0.188), a neurotransmitter involved in gut-brain signaling and motility. **Group Definitions:** No exposure = no glyphosate exposure in F0; AAD = Average American Diet intake level (0.01 mg/kg/day); EPA Upper Limit = U.S. EPA acceptable daily intake (1.75 mg/kg/day.) “F1” and “F2” refer to first- and second-generation offspring of F0-exposed breeders. Genotypes: *Muc2*^+/-^ = healthy mice; *Muc2*^-/-^ = colitis-susceptible mice. **Sex:** Sex was not found to be a significant predictor for changes observed in distance (PERMANOVA, *p*=0.757,), velocity (PERMANOVA, *p*=0.792), total rearing events (PERMANOVA, *p*=0.173), unsupported rearing events (PERMANOVA, *p*=0.167), during radial-arm maze test (PERMANOVA, *p*=0.964) or serum kynurenine levels (PERMANOVA, *p*=0.346) in healthy mice. Sex was not found to be a significant predictor for changes observed in serum serotonin levels (PERMANOVA, *p*=0.662), alpha-synuclein (PERMANOVA, *p*=0.856), SNAP-25 (PERMANOVA, *p*=0.212) or GFAP expression (PERMANOVA, *p*=0.195). **Statistical Summary:** Confidence intervals, and sample sizes broken down by sex provided in table S6.

While colitis-susceptible (*Muc2*^-/-^) offspring did not show behavioral impairments, F2 AAD and EPA groups exhibited increased colonic expression of α-synuclein, GFAP, and SNAP-25 (*p* < 0.05, FDR-corrected; Fig. 3E), consistent with enteric neuroinflammation and early neurodegenerative signaling. EPA F2 *Muc2*^-/-^ mice also had lower serum serotonin (*p* = 0.031, η² = 0.188; Fig. 3F), a key neurotransmitter involved in gut-brain signaling and motility. Although behavioural phenotypes were restricted to healthy offspring, the gut-brain molecular changes observed in colitis-susceptible mice suggest latent neurophysiological disruption. Together, these findings show that prenatal glyphosate exposure, at or below levels deemed safe, can affect neurodevelopment and behaviour through gut-brain axis pathways.

### Prenatal glyphosate exposure alters gut bacteriome composition, microbial metabolism, and predicted functional potential in offspring

While community structure was not significantly altered when examining alpha- and beta-diversity metrics, prenatal glyphosate exposure induced taxonomic shifts in both healthy (*Muc2*^+/-^) and colitis-susceptible (*Muc2*^-/-^) offspring across generations, with genotype-specific patterns emerging (Fig 4). In healthy offspring, *Akkermansia muciniphila* was more abundant in unexposed controls. Given its role in supporting barrier integrity and regulating GLP-1^26^ and NF-κB signaling,^27^ its depletion in exposed groups may underlie both metabolic and inflammatory phenotypes.

**Figure 4.**
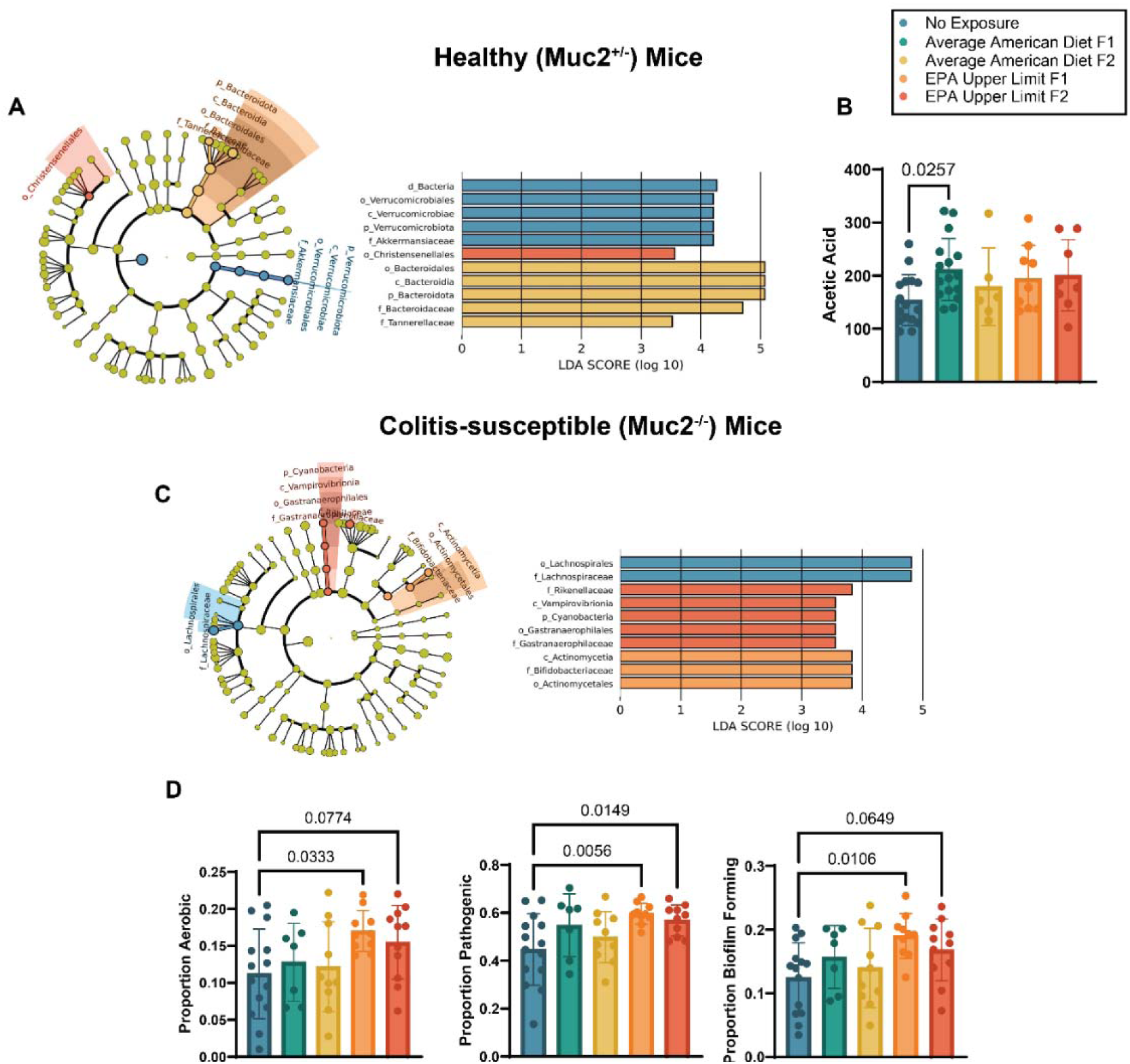
Prenatal glyphosate exposure alters gut microbial composition, SCFA levels, and predicted microbial function across generations. **(A)** Microbial taxa affected by prenatal glyphosate exposure in healthy (*Muc2*^+/-^) offspring included *Parabacteroides distasonis*, *P. goldsteinii*, and Christensenellaceae, all of which were significantly elevated in AAD or EPA F2 mice. Notably, *Akkermansia muciniphila* was more abundant in unexposed offspring, consistent with its role in maintaining gut barrier integrity and maintaining metabolic homeostasis. (B) Cecal acetate levels were significantly increased in AAD F1 offspring (*p* = 0.064, η²p = 0.144), suggesting altered microbial fermentation and potential implications for host metabolic regulation. (C) In colitis-susceptible (*Muc2*^-/-^) mice, glyphosate exposure was associated with enrichment of *Bifidobacterium* spp. (EPA F1) and Gastranaerophilales (EPA F2), including non-photosynthetic cyanobacteria suggesting a possible interaction between glyphosate and microbial phosphorus metabolism. (D) BugBase functional prediction revealed that *Muc2*^-/-^ offspring exposed to EPA doses harbored microbiota with increased proportions of aerobic (*p* = 0.0676, η²p = 0.167), biofilm-producing (*p* = 0.0365, η²p = 0.192), and pathogenic bacteria (Welch’s ANOVA, *p* = 0.0137, ω² = 0.370), consistent with a dysbiotic microbial community. Group Definitions: No exposure = no glyphosate exposure in F0; AAD = Average American Diet intake level (0.01 mg/kg/day); EPA Upper Limit = U.S. EPA acceptable daily intake (1.75 mg/kg/day.) “F1” and “F2” refer to first- and second-generation offspring of F0-exposed breeders. Genotypes: *Muc2*^+/-^ = healthy mice; *Muc2*^-/-^ = colitis-susceptible mice. Sex: Sex was not found to be a significant predictor of acetic acid production in healthy mice (PERMANOVA, *p*=0.739) nor any of the predicted functional analysis conducted in colitis-susceptible mice (PERMANOVA, aerobic *p*=0.947; pathogenic *p*=0.873, biofilm *p*=0.991). Statistical Summary: Confidence intervals, and sample sizes broken down by sex provided in table S6.

In contrast*, Parabacteroides distasonis* and *P. goldstein* were elevated in AAD F2 mice. While sometimes considered beneficial, *P. distasonis* has been linked to depression-like behavior in murine colitis models, pointing to context-dependent effects on host physiology.^28^ EPA F2 mice also showed increased Christensenellaceae, a highly heritable taxon associated with gut-brain signaling and Parkinson’s disease risk.^29,30^ These changes suggest glyphosate exposure shifts the microbial ecosystem toward profiles linked to metabolic and neurological vulnerability. Cecal metabolite analysis revealed elevated acetate in AAD F1 mice (Fig. 4B) While short-chain fatty acids are typically beneficial, excess acetate has been associated with parasympathetic overactivation and features of metabolic syndrome, suggesting a potential mechanism by which even low-dose glyphosate might disrupt host energy balance.^31^

In colitis-susceptible (*Muc2*^-/-^) offspring, *Bifidobacterium* spp. were increased in EPA F1 mice. Although typically regarded as beneficial, elevated *Bifidobacterium* spp. have been reported during active IBD flares, suggesting its expansion may reflect intestinal inflammation.^32^ Additionally, colitis-susceptible (*Muc2*^-/-^) F2 offspring within the EPA Upper Limit group exhibited an increased abundance of non-photosynthetic cyanobacteria, specifically Gastranaerophilales. In the environment, glyphosate promotes the growth of cyanobacteria as it provides a rich source of phosphorus needed for these organisms to thrive.^33^ However, this study is the first to demonstrate that dietary glyphosate can promote the growth of cyanobacteria in the gut. Cyanobacteria within the environment can produce the non-coding amino acid β-Methylamino-L-alanine (BMAA). BMAA can be incorporated into proteins where it acts as a neurotoxin through the misfolding of proteins and has been associated with neurodegenerative diseases like Parkinson’s. It has been suggested that glyphosate runoff may contribute to algae blooms, increasing BMAA levels and contributing to the recent increase in neurological diseases observed in coastal areas across North America.^34,35^

Functional profiling using BugBase^36^ showed colitis-susceptible EPA offspring harbored a bacteriome enriched for aerobic, pathogenic, and biofilm-forming taxa across both F1 and F2 generations (Fig. 4D.) These features are consistent with microbial dysbiosis and may exacerbate barrier disruption and immune activation. Given that glyphosate targets the EPSPS enzyme in the Shikimate pathway—and that most commensals possess Class I EPSPS enzymes (glyphosate-sensitive), while several pathogens carry resistant Class II variants, these shifts likely reflect a selective pressure favoring glyphosate-tolerant taxa.^7,8^ These results show that prenatal glyphosate exposure reconfigures the gut microbiome in ways that may promote inflammation, metabolic dysfunction, and neuroimmune disruption. The persistence of these shifts across generations and their emergence at human-relevant doses highlights their potential significance for long-term health.

## Discussion

The findings of this exploratory study show that prenatal glyphosate exposure, even at doses previously considered safe, induces measurable physiological changes in offspring across generations. In healthy (*Muc2*^+/-^) mice, glyphosate exposure induced colonic histopathology marked by goblet cell depletion and epithelial hyperplasia. Loss of goblet cells reduced mucin-2 production and weakened the mucus barrier, which may have allowed microbial translocation and contributed to immune activation, as indicated by elevated serum LIX. These effects were absent in colitis-susceptible (*Muc2*^-/-^) mice, suggesting that glyphosate primarily impacts barrier integrity in hosts with intact immune and mucus signaling.

Both genotypes showed a shift toward a pro-inflammatory TH1/TH17 cytokine profile. These findings contrast with asthma model studies where glyphosate suppressed IL-17 and IFN-γ, highlighting how cytokine responses may vary by tissue and context.^37^ Glyphosate’s known non-monotonic dose-response behaviour may also contribute, where effects emerge at low or high doses but are not always linear.^38^ All animals received trace glyphosate from chow (∼0.0015 mg/day), but the additional waterborne exposure in AAD and EPA groups was enough to drive phenotypes, especially at the lower AAD dose. These findings support a threshold-based, non-monotonic model of toxicity. Several effects, including working memory deficits and acetate accumulation, were observed in AAD but not EPA animals, underscoring the need to rethink dose-response assumptions for environmental exposures.

Metabolic disruption was evident in healthy (*Muc2*^+/-^) AAD and EPA offspring, with reduced GLP-1 and altered leptin and ghrelin levels contributing to impaired glucose tolerance and insulin resistance. These findings align with CHAMACOS cohort data linking early-life glyphosate exposure to increased metabolic syndrome risk by age 18. Interestingly, later-life urinary glyphosate or AMPA levels were not predictive, reinforcing the idea that there is a specific developmental window of susceptibility to the toxic effects of glyphosate.^39^

Behaviorally, healthy (*Muc2*^+/-^) EPA-exposed F2 mice showed reduced locomotor and exploratory activity and lower serum kynurenine—a precursor to neuroactive metabolites like kynurenic and quinolinic acid. An imbalance in this pathway can increase vulnerability to excitotoxicity and neuroinflammation, potentially contributing to observed behavioral impairments.^40^ In contrast, colitis-susceptible (*Muc2*^-/-^) offspring did not exhibit behavioral changes but did show signs of enteric neuroinflammation. SNAP-25, GFAP, and α-synuclein were highly expressed in AAD and EPA F2 groups, suggesting enteric neuronal stress and glial activation.^41-43^ These changes are consistent with gut-brain signaling disruption and may represent early neurodegenerative processes which have been observed following acute high-dose glyphosate exposure.^44-46^

Microbiome shifts were observed across genotypes and generations. In healthy (*Muc2*^+/-^) offspring, *Akkermansia muciniphila* was depleted, while *Parabacteroides* spp. and Christensenellaceae were enriched. Though sometimes considered beneficial, these taxa have been linked to behavioral and metabolic risk in context-specific ways. In colitis-susceptible (*Muc2*^-/-^) offspring, *Bifidobacterium* spp. was elevated in EPA F1 mice, similar to what is observed during active IBD in humans. Most striking was the increased abundance of Gastranaerophilales in EPA F2 mice. These cyanobacteria are potential BMAA producers, raising the possibility of gut-derived neurotoxic exposure. Glyphosate’s phosphonate group may select for microbes with enhanced phosphorus acquisition, offering one explanation for the emergence of this rare lineage. Importantly, *Muc2*^+/-^ and *Muc2*^-/-^ littermates were co-housed to minimize cage effects and standardize early microbial exposures. While co-housing enables microbial exchange, it reduces inter-cage variation and strengthens causal inference around treatment effects.^47^ BugBase^36^ profiling supported a shift toward dysbiosis in colitis-susceptible (*Muc2*^-/-^) EPA offspring, with increased aerobic,^48^ pathogenic,^49^ and biofilm-forming^50,51^ taxa— features associated with inflammation and barrier breakdown.

F0 breeders were not tested metabolically or behaviorally to avoid stress-induced confounds, so we cannot distinguish between inherited epigenetic changes and microbiota-mediated effects. However, the appearance of phenotypes in the unexposed F2 generation suggests transgenerational transmission, likely involving both host and microbial pathways. However, future gnotobiotic studies are needed to test these mechanisms and show a causal relationship.

These findings show that prenatal glyphosate exposure, at doses consistent with real-world intake, can disrupt multiple physiological systems across generations. The emergence of effects at low doses highlights the potential for non-monotonicity to mask toxicity in traditional testing frameworks. Rethinking regulatory thresholds and expanding mechanistic studies, especially those focused on developmental timing, microbiome involvement, and intergenerational transmission, will be critical to thoroughly assessing glyphosate’s long-term health impact.

## Materials and Methods

### Pre-registration

The original experimental design was pre-registered with the Open Science Framework (osf.io) on February 25th, 2020. The pre-registration be viewed at osf.io/6nbsx. Changes made from original pre-registration can be found in table S1.

### Determination of the Average American Diet (AAD) dose

To determine the amount of glyphosate the average North American is exposed to through diet, we calculated the Average American diet (AAD) dose based on a hypothetical menu for a 60kg female following the American food guide and reported values of glyphosate found within these foods (Table 1.) The AAD dose was determined to be 0.01mg/kg/day, which is more than 100X lower than the acceptable daily intake (ADI) currently set by the Environmental Protection Agency (EPA) at 1.75 mg/kg/day. At the time of dose calculation, few studies existed detailing glyphosate levels in food. Therefore we relied on values collected from citizen science initiatives to calculate the AAD dose. To validate these reported levels of glyphosate in food, we developed a modified ultra-performance liquid chromatography method for quantification (figure S2.) The values used to calculate the AAD dose were found to be similar to what has subsequently been reported in literature and our own measurements (Table 2.)

**Table 1:** Calculation of the AAD dose. A hypothetical menu for an active female > 20 years old (60kg) following the American food guide.

| Meal | Amount | Glyphosate Detected (ppb) | Total (µg/serving) |
| --- | --- | --- | --- |
| **Breakfast** |  |  |  |
| Cheerios | 1 cup (27g) | 5775<sup>2</sup> | 15.6 |
| 1% Milk | 1 cup | - | - |
| Apple, whole | 1 medium | - | - |
| **Snack** |  |  |  |
| Ritz crackers | 10 (32g) | 569<sup>52</sup> | 18.2 |
| Fontaine Sante roasted garlic hummus | 1/4 cup (60g) | 760<sup>52</sup> | 45.6 |
| Water | No limit | - | - |
| **Lunch** |  |  |  |
| PC Blue Menu Tortilla (100% whole grain) | 2 (85g) | 744<sup>52</sup> | 63.2 |
| Salsa | 2 tbsp | - | - |
| Ground beef (grain fed and finished) | 1/2 cup (75g) | 5,000*<sup>53</sup> | 375 |
| Mozzarella cheese | 2 ounces | - | - |
| Carrot/celery sticks | 1 cup | - | - |
| **Snack** |  |  |  |
| Tim Hortons glazed Timbit | 6 (150g-25g each) | 209<sup>52</sup> | 31.4 |
| Decaf coffee | 1 cup | - | - |
| Honey | 1 tbsp (21g) | 64<sup>54</sup> | 1.34 |
| 365 coffee cream | 2 tbsp (30g) | 104<sup>55</sup> | 3.12 |
| **Dinner** |  |  |  |
| Grilled salmon | 4 oz | - | - |
| Lentil salad | 1/2 cup (80g) | 284<sup>56</sup> | 22.7 |
| Green beans | 1 cup | - | - |
| Teriyaki sauce (with soy) | 2 tsp (8g) | 242<sup>57</sup> | 1.93 |
| Apple crisp made with oatmeal | 1/2 cup (50g) | 135<sup>52</sup> | 6.75 |
| Water | No limit | - | - |
| **Snack** |  |  |  |
| Ritz crackers | 10 (32g) | 5695<sup>2</sup> | 18.2 |
| Cream cheese | 2 tbsp | - | - |
| Apple, sliced | 1/2 medium | - | - |
| **Total (µg/day)** |  |  | **603.02** |
| **Total (mg/kg/day)** |  |  | **0.01** |

\*Maximum Allowable Amount

**Table 2:** Glyphosate levels in commercially available foods and laboratory rodent chow using our modified UPLC detection method.

| Sample Matrix | Concentration of Glyphosate<br>(µg/g) | Concentration<br>Glyphosate (ppb) |
| --- | --- | --- |
| Red Lentils | 0.245 | 245 |
| Picolab® 5053 Mouse Diet | 0.381 | 381 |
| Desiccated Wheat | 2.445 | 2445 |
| Nondesiccated Wheat | ND | ND |

We also assessed glyphosate content in the irradiated PicoLab® 5053 rodent chow used in this study. Tandem MS analysis revealed a glyphosate concentration of 0.381 μg/g (381 ppb.) Based on typical chow consumption (4–5 g/day),^58^ this corresponds to a total glyphosate intake of approximately 0.00152–0.00191 mg/day per mouse. This background exposure was not adjusted for body weight and was consistent across all experimental groups, including non-exposed controls. Therefore, animals in the AAD and EPA groups received this baseline amount plus their respective water-based doses of 0.01 mg/kg/day and 1.75 mg/kg/day, respectively. While glyphosate-free chow was not commercially available at the time of this study, this background exposure was minimal and equivalent across groups, allowing differences between AAD, EPA, and non-exposed control animals to be attributed to the additional, controlled waterborne exposure.

### Animal husbandry

Mice were housed under specific pathogen-free conditions within the bioscience facility at UBC Okanagan. The UBC Animal Care Committee approved all animal experiments under protocols A22-0231 and A18-0317. Mice were kept on ventilated racks and housed at 22 ± 2°C with controlled humidity on a 14h:10h light: dark cycle and provided with autoclaved woodchips, crinkle paper, and polycarbonate houses for cage enrichment. Parental generation *Muc2*^+/-^ and *Muc2*^-/-^ mice were bred in-house. Original *Muc2*^-/-^ mice used to generate our colony were obtained from Dr. Bruce Vallance at UBC Vancouver and were derived on a C57Bl/6 background. To mitigate cage effects, we bred heterozygous (*Muc2*^+/-^) with homozygous (*Muc2*^-/-^) mice, thereby creating co-housed littermates who were healthy (*Muc2*^+/-^) and colitis-susceptible (*Muc2*^-/-^.) This has been shown to be an optimal method for microbiome standardization. Within our facility, we have shown that *Muc2*^+/-^ are healthy and do not develop colitis or any metabolic or behavioral abnormalities associated with a total knockout of *Muc2* and therefore, served as our healthy population. A comparison of *Muc2^+/+^, Muc2^+/-^* and *Muc2^-/-^* mice within our facility can be found in (figure S1, A-D.)

### Sample size justification

Given the unique animal model, breeding scheme, and proprietary dose used in this study, it was impossible to collect the effect size and standard deviation from literature. Therefore, we estimated sample size using the resource equation approach.^59-63^ In brief, the resource equation is based on the law of diminishing returns. It assumes that the acceptable range of degrees of freedom (DF) for the error term within an analysis of variance (ANOVA) is between 10 and 20. For one-way ANOVA, the within-subject DF can be determined as:

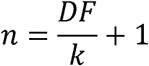

Where:

is the number of groups
is the number of subjects per group.

Based on the acceptable range of the DF, the DF in the formula is replaced with the minimum (10) and maximum (20) to obtain the minimum and maximum numbers of animals per group. Therefore, the minimum and maximum number of animals needed for the current study was estimated for k = 5 groups (No Exposure, Average American Diet F1, Average American Diet F2, EPA Upper Limit F1 and EPA Upper Limit F2.) Using the equation above, it was determined that each group requires a minimum of three animals and a “maximum” of five per genotype. However, it should be noted that the resource equation is not as robust as the power analysis method, and every effort was made to include more than the maximum number of animals required per group. A post hoc power analysis was not conducted given that probability refers to the likelihood of a future event occurring; therefore, conducting a power analysis when the known discovery of a significant finding has occurred is not appropriate and potentially misleading.^65^

### Experimental design

**Fig. 5:**
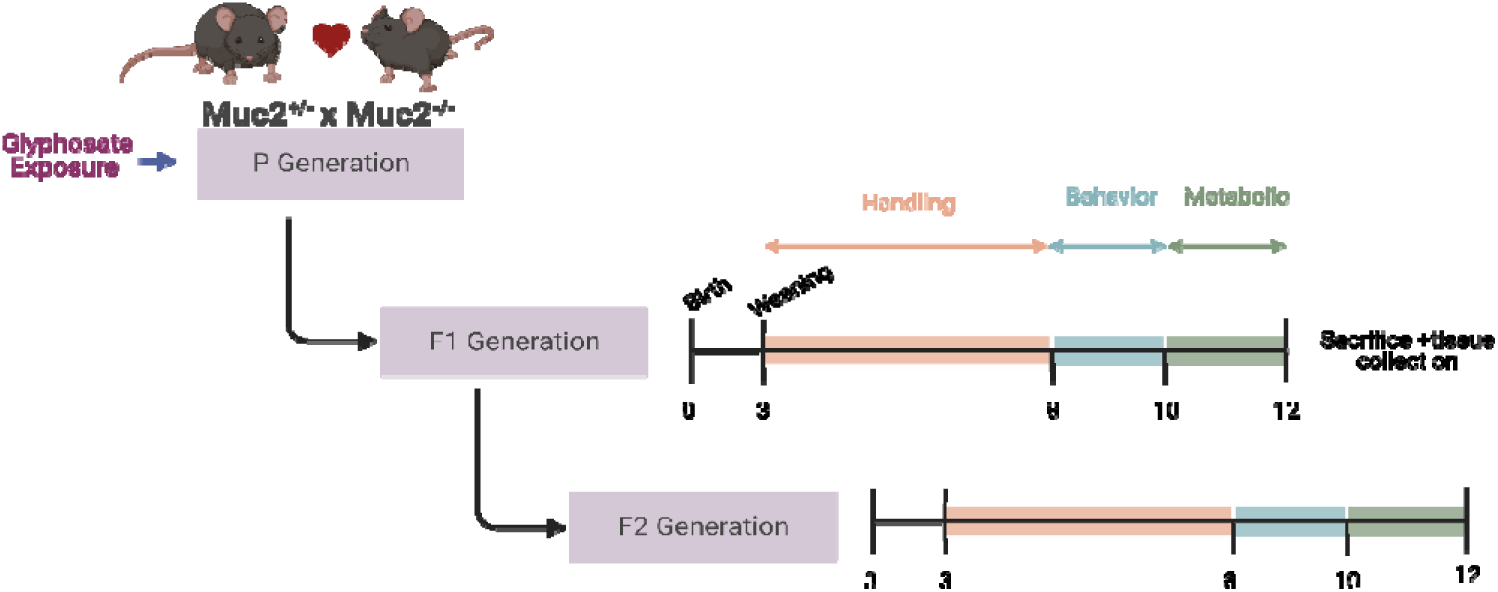
Experimental design.

Upon reaching sexual maturity (6 weeks of age), breeding pairs consisting of *Muc2*^+/-^ (healthy) and *Muc2*^-/-^ (colitis susceptible) mice. A summary of parental generation sex and genotype combinations can be found in table S2. Non-exposed mice received autoclaved reverse-osmosis water *ad libitum*. F0 breeders began receiving glyphosate-supplemented water at sexual maturity and exposure continued through mating, gestation. Glyphosate-exposed mice received autoclaved reverse-osmosis water with glyphosate N-(phosphonomethyl)glycine 96%, Millipore Sigma, Cat ID: 337757) at either the AAD dose (0.01mg/kg body weight/day) or the current ADI set by the EPA (1.75mg/kg body weight/day), *ad libitum*. Non-exposed animals were generated in parallel with exposed cohorts and tested within the same experimental windows to minimize batch effects across generations. Researchers were not blinded with regards to treatment, however experimenters were blind with regards to genotype of animals within the cage (genotyping occurred at the end of the experiment as to not influence handlers. All mice received irradiated PicoLab® Mouse Diet 5058 *ad libitum*. At six weeks of age, offspring were assigned using a random number generator to one of two fates - experimental or breeding. Care was taken to ensure that random assignment occurred with no sibling or cousin breeding to avoid inbreeding. Breeding mice were used to generate the F2 generation, whereas experimental mice were handled weekly until seven weeks of age and then daily for one week before behavior testing began. Mice in the experimental group began behavior testing at eight weeks and metabolic testing at ten weeks. Mice were sacrificed at 12 weeks of age. In brief, mice were anesthetized to the surgical plane using isoflurane, and cardiac blood was collected via cardiac puncture. The time between cage disturbance and death was less than three minutes, as exceeding that time elicits stress hormone production.^66^ Death was confirmed using cervical dislocation, and the distal colon, ileum, stool, cecum, and liver were collected for analysis. Both the distal colon and ileum were divided into three pieces, with one piece being flash-frozen in liquid nitrogen and stored at -80°C for microbiome analysis, one stored in RNAprotect (Qiagen, Cat ID: 76106) for mRNA analysis, and the third in 10% neutral buffered formalin (Fisher Scientific, Cat ID: 23-305510) for histopathological analysis. The collected cardiac blood was spun for 15 minutes at 1,500 g to isolate the serum. Once F2 generation pups were weaned, breeding mice were sacrificed, and tissues were collected as outlined previously. Each mouse was considered to be an experimental unit. No metabolic or behavioral phenotyping was conducted on F0 breeders to avoid introducing stress-related variability or confounds during mating, gestation, and lactation. All outcomes were assessed exclusively in F1 and F2 offspring to isolate effects of prenatal exposure and potential intergenerational transmission.

### Open field test

At eight weeks of age, mice were habituated to the testing room for one hour before test initiation. The open field setup comprised a single 40 cm x 40 cm x 40 cm arena made of opaque white, non-reflective acrylic material (Conduct Science). The box was cleaned with 10% ethanol before and between sessions, reducing stress odors and olfactory cues. A video camera (GoPro Hero 8) recorded the mouse activity within the open field. Mice were placed in the center of the open field, and their activity was recorded for ten minutes. The testers positioned themselves outside the mice’s field of view, and mice were monitored remotely using the GoPro Quik App (version 12.4.1, Apple iPhone iOS). Video footage was analyzed using EthoVision XT (version 17; Noldus) for the number of center square entries, time spent within the center square, distance travelled, body velocity, rearing frequency, and time spent grooming. Data was collected from two independent experiments.

### Radial arm maze

Mice were assessed using the fully baited training procedure where each arm chamber contained a food reward, and the mouse was expected to learn to visit each arm only once per session. Once the mouse has obtained the food reward from an arm, re-entry of the now unabated arm is considered a memory error. The radial arm maze test was performed over four consecutive days. Visual cues outside the maze remained constant over the four days to allow the mice to establish a mental map of the maze. As mice are neophobic, treats were provided to mice in their home cages three days before testing began. **Day 1:** The mice were habituated to the maze and allowed to explore it freely for ten minutes. **Days 2-4:** Mice were placed in the center of the radial arm maze until task completion was achieved. Task completion was defined as entry into each arm of the maze or ten minutes had elapsed. Mice were monitored remotely to determine the number of times an arm had been entered (memory errors). The total number of memory errors was averaged over three consecutive trials.

### Light/dark test

Forty-eight hours after open-field testing, the mice were again habituated to the testing room for one hour before test initiation. The light/dark setup consisted of a single 40 cm x 40 cm box divided into two compartments, with one being open to the light and the other being dark (light compartment: 25 cm x 40 cm, dark compartment: 17.5 cm x 40 cm) (Conduct Science). Mice were placed in the center of the light section of the light/dark maze, and their activity was recorded for ten minutes (GoPro Hero 8). The observers positioned themselves outside the mouse’s view, and the maze was wiped with 10% ethanol before each session and between sessions, as described above. An impartial, blinded individual manually analyzed the video footage. Metrics measured included the time spent in the dark, the number of transitions between light and dark, and the latency to enter the dark compartment.

### Oral glucose tolerance test

Mice were fasted for six hours before the start of the test which has been shown to be optimal in rodents.^67^ Fasting blood glucose readings were obtained from blood samples collected from a tail poke and quantified using a OneTouch Verio Flex® glucometer. Before testing, the glucometer was calibrated using the control solution provided by the manufacturer. A 30% glucose solution was prepared using D-(+)-Glucose (Sigma Aldrich, Cat ID: 50-99-7) prepared in phosphate-buffered saline and sterile-filtered using a 0.2-micron syringe filter. Following the fasting blood glucose reading, mice received glucose gavage at a dose equal to 2g/kg fasted body weight. Blood glucose readings were collected 15-, 30-, 60-, and 120 minutes after gavage. The area under the curve (AUC) was determined using the trapezoid rule.

### Insulin tolerance test

Mice were fasted for six hours before the start of the test. Initial blood glucose readings were obtained as described above. Human insulin (Humalog®, 100U/ml) was diluted to a 0.1U/ml working solution by sterile isotonic saline. Following the initial blood glucose reading, the mice received an intraperitoneal insulin injection dose equal to 0.5U/kg fasted body weight. Blood glucose readings were collected 15-, 30-, 60-, and 120 minutes after injection. If at any time a mouse became hypoglycemic, defined as a blood glucose reading less than 2.3mmol/L, or if blood glucose levels failed to return to normal following 120 minutes, mice received a bolus intraperitoneal injection of a 30% sucrose solution (Sigma Aldrich, Cat ID: 57-50-1) and were removed for the remainder of the test. AUC was determined using the trapezoid rule.

### Histopathological scoring

Tissues collected for histology were immediately placed in 10% neutral buffered formalin. Following 24-hour formalin fixation, the tissues were washed in PBS and stored at 4°C in 70% ethanol. Tissues were sent to BC Children’s Hospital Research Institute for paraffin embedding, sectioning, and hematoxylin and eosin (H&E) staining. Coded samples blinded to the scorers were evaluated and scored by three people, and scores were averaged. Samples were collected from two independent experiments. Scoring parameters used can be found in table S3 and S4.

### Immunohistochemistry

Tissues were collected as outlined above. Unstained paraffin slides were cut for immunohistochemistry. Slides were deparaffinized in xylene (two five-minute washes), and tissues were rehydrated using a series of ethanol washes ranging from 100% - to 70% ethanol before being rinsed in deionized water and phosphate-buffered saline (PBS.) Tissues were then incubated with trypsin (Sigma Aldrich Cat-ID: 9002-07-7) at 37°C for 30 minutes to facilitate antigen retrieval. Following antigen retrieval, a blocking solution of 5% IgG-free bovine serum albumin (BSA) (Sigma Aldrich Cat-ID: 9048-46-8) was applied, and slides were incubated for 20 minutes at room temperature. A primary antibody solution (F4/80: Cedarlane, Cat-ID: CL89170AP, MPO: Invitrogen, Cat-ID: PA516672, ZO2: Invitrogen, Cat-ID: 71-1400) was applied at a 1:50 antibody: BSA dilution ratio and slides were incubated overnight at 4°C. After incubation, slides were rinsed in PBS. A fluorescently conjugated secondary antibody (F4/80: Invitrogen, Cat-ID: A11006, MPO: Invitrogen, Cat-ID: A11012, ZO2: Invitrogen, Cat-ID: A-11008) was applied at a 1:1000 antibody: BSA dilution ratio and incubated at room temperature for one hour. Following incubation, a histology mounting medium containing 4’-6-diamidino-2-phenylindole (DAPI) was applied (Sigma Aldrich Cat-ID: F6182) and a coverslip was applied. Slides were imaged using the EVOS M500 cell imaging system (ThermoFisher Cat-ID: AMF5000SV). Cells were quantified using QuPath (version 0.4.4) using positive cell detection to ensure cells counted colocalized both fluorescent signals (DAPI stained nucleus and fluorescently stained structure of interest).

### RNA Sample preparation

According to the manufacturer’s instructions, total RNA was extracted from the colon tissue using the RNeasy Fibrous Tissue Mini Kit (Qiagen, Cat ID: 74704). 2-mercaptoethanol (Sigma Aldrich, Cat ID: M6250) was added to the lysis buffer to protect against RNase degradation. All kits were ordered simultaneously, and samples were randomized across kits to minimize potential batch effects. Immediately following extraction, purified RNA was quantified spectrophotometrically (ThermoFisher, Nanodrop 2000), normalized to 250ng/μl and cDNA was synthesized using an iScript Reverse Transcription kit (BioRad, Cat ID: 1708890) according to the manufacturer’s instructions with 500ng RNA per reaction. cDNA was diluted to a final concentration of 5ng/μl for downstream analysis. Samples were collected from two independent experiments.

### Quantitative real-time PCR

Relative gene expression was quantified using quantitative real-time PCR (qPCR). The 10μl reaction consisted of 0.4μl of each forward and reverse primer (10mM), 5μl SsoAdvanced Universal SYBR Green Supermix (BioRad, Cat ID: 1725210), 3.2μl DNase-free water and 1μl cDNA template. Samples were assigned to a plate using a random number generator to mitigate batch effects. Samples were run in duplicate using primers separated by at least one intron on the corresponding genomic DNA to prevent amplification of genomic DNA when possible; for primers not separated by one intron, controls containing no reverse transcriptase were run to ensure that there was no amplification of genomic DNA. A list of primer sequences can be found in table S5.

### Relative gene expression

Relative gene expression was determined using Bio-Rad CFX Manager software (version 2.1, Hercules, California, USA). The relative normalized expression was determined by dividing the relative expression of the gene of interest by the geometric mean of the relative expression of the reference genes. The log of the relative normalized expression is equivalent to the ΔΔCq. The reference genes *Eef2* and *Tbp* were used as they are stable within the gut during inflammation.^68^ Primer efficiencies for each assay were determined using LinRegPCR (version 2021.2), which uses the slope of the exponential phase of each reaction to determine the least variable mean PCR efficiency per assay. This is superior to the standard curve method as it allows for a more precise determination of amplification efficiency and is more sensitive in detecting slight differences in the initial amount of template. Additionally, LinReg can provide reliable quantification in the presence of minor experimental variations (e.g., reaction efficiency, primer-dimer formation). qPCR data were generated using a BioRadC1000 Touch Thermal Cycler (CFX96) and analyzed using Bio-Rad CFX Manager Software version 2.1 (Hercules, California, USA). Quality control measures were conducted on each run, including negative control with a Cq less than 38, primer efficiency below 90% or over 110%, and replicate group standard deviation greater than 0.20.

### Cytokine network analysis

Cytokine network analysis was conducted in R Studio (version 2023.09.01 Build 494) running R (version 4.3.1) using packages *tidyverse* and *factoextra*.

### Metabolic hormone analysis

Blood was collected from mice via cardiac puncture. The serum was isolated by allowing the blood to clot at room temperature for 20 minutes before spinning at 1,500 g for 15 minutes. A protease inhibitor cocktail (Amresco, Cat ID: M221) was added to sera before being stored at - 80°C for analysis. Serum samples that were hemolyzed or lipemic were not used. Serum samples were diluted 2-fold with sterile PBS (pH=7.5). Serum was sent to Eve Technologies (Calgary, AB, Canada) and analyzed for metabolic hormones (Mouse Metabolic Array) using addressable laser bead immunoassay.

### Enzyme-linked immunosorbent assay (ELISA) analysis

Serum corticosterone levels were quantified using the 96-well Corticosterone Multi-Format ELISA kit (Arbor Assays, Cat ID: K014-H5.) Serum samples were collected, aliquoted, and stored at -80°C before analysis. Serum samples that were hemolyzed or lipemic were not used. All samples were diluted 100-fold before use. This kit is sensitive to 20.9pg/mL using 50μl input. All samples were randomly assigned to a plate using random number generation and were run in duplicate according to the manufacturer’s specifications. Plates were read at 450nm, and baseline optical density correction was conducted at 620nm. Concentrations were determined using MyAssays (MyAssays.com.)

### Microbiome Sample preparation

Total microbial DNA was isolated from colon tissues using the QIAmp PowerFecal Pro DNA Kit (Qiagen, Cat ID: 51804.) All kits were ordered simultaneously, and samples were randomized across kits to minimize potential batch effects. In brief, samples were added to the PowerBead Pro Tube™ and homogenized for 20 minutes using a vortex adapter (Qiagen, Cat ID:13000-V1-24.) The rest of the extraction was conducted according to the manufacturer’s specifications, except for an additional wash with the included ethanol-based wash solution to improve sample purity. DNA was quantified spectrophotometrically (ThermoFisher, Nanodrop 2000). A mock community was also extracted along with samples and sent for sequencing. Integrated Microbiome Platforms provided the mock community for Advancing Causation Testing and Translation (IMPACTT) (Calgary, AB, Canada.) Samples were collected from two independent experiments.

### 16s sequencing

DNA samples were sent to Gut4Health (BC Children’s Hospital Research Institute, Vancouver, BC) for library prep and 16s sequencing. In brief, the V4 region of the 16s gene was amplified with barcode primers containing the index sequences using a KAPA HiFi HotStart Real-time PCR Master Mix (Roche.) PCR product amplification and concentration were monitored using a Bio-Rad CFX Connect Real-Time PCR system. Amplicon libraries were then purified, normalized, and pooled using the SequalPrep™ normalization plate (Applied Biosystems.) According to the manufacturer’s instructions, library concentrations were verified using a Qubit™ dsDNA high-sensitivity assay (Invitrogen) and KAPA Library Quantification Kit (Roche.) The purified pooled libraries were submitted to the Bioinformatics and Sequencing consortium at UBC Vancouver, which verified DNA quality and quantity using an Agilent high-sensitivity DNA kit on an Agilent 2100 Bioanalyzer. Sequencing was performed on the Illumina MiSeq™v2 platform with 2 x 250 paired-end read chemistry.

### Bioinformatics

All bioinformatic processes were performed using a combination of R statistical software and QIIME 2 (version 2023.9) using the various built-in plugins described below. Demultiplexed sequences obtained from the sequencing facility underwent quality control and denoising using DADA2. Forward and reverse reads were truncated at 240 bases. Taxonomy was assigned to sequences using a pre-trained Naïve Bayes classifier trained on the Greengenes version 2 classifier (version 2022.10.backbone.v4.) Differential abundance testing and effect size of taxa abundances were conducted using LEfSe, where features with fewer than ten counts across all samples or appearing in fewer than five samples were removed. A multiclass comparison was performed using a Kruskal-Wallis test (α=0.05), and a linear discriminant analysis (LDA) score for discriminative features was set to 2.0. Biologically relevant features were visualized in a cladogram. BugBase was used to predict high-level community phenotypes present within samples. For BugBase analysis, VSEARCH was used to filter the feature table generated by DADA2 using a closed-reference approach. Samples were filtered using the Greengenes reference database (version 13.8) and 97% clustering.

### Statistical analysis

All statistical analysis was conducted in GraphPad Prism (version 10) unless otherwise specified. Data displaying a log-normal distribution was log-transformed before outlier and statistical analysis and is displayed in its log-transformed version. Outliers were detected using the ROUT method with Q=1%. Statistical assumptions, including normality and homogeneity of variance, were tested using D’Agostino-Pearson omnibus (K2) and Bartlett’s test, respectively, and these tests were used to determine which statistical test was used. Multiple comparisons were corrected by controling the false discovery rate using the two-stage step-up method of Benjamini, Krieger and Yekutieli. For parametric tests, effect size was calculated using the method described by Keppel and Wickens and is reported as partial eta squared (η^2^p).^69^ For non-parametric tests, eta-squared (η²) was calculated manually using the formula:

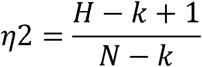

where:

*H* is the Kruskal-Wallis test statistic.
*k* is the number of groups.
*N* is the total number of observations across all groups.

A result was considered significant with an adjusted *p*-value ≤ 0.05. Results with an adjusted *p*-value ≤ 0.08 were considered a non-significant trend. Confidence intervals (CI) reported represent the 95% confidence interval. Due to the inherent unpredictability of mouse breeding, we were unable to obtain sufficient numbers of each sex within each generation and genotype to achieve the statistical power needed for a detailed analysis of sex differences. To address this limitation, we conducted an exploratory analysis of potential sex effects using a permutational multivariate analysis of variance (PERMANOVA) with 999 permutations, implemented via the *adonis2* function in the *vegan* R package. This analysis assessed the influence of group, sex, and group × sex interactions, using Euclidean distance metrics. The reported PERMANOVA *p*-values represent the combined effects of group and sex.

## Data and materials availability

The 16S sequences files and metadata used for the analysis of this study are publicly available at: https://doi.org/10.5061/dryad.ht76hdrqj.70 [currently under review, peer-review link found here: https://datadryad.org/stash/share/O0oWxMXzf-eLgBZv31_ZF7eC6iXXbWXn5URRInd2GGo]

## Acknowledgements

We would like to thank our funders, including crowdfunding UBC, the Natural Sciences and Engineering Research Council (NSERC) who supported this work through both a discovery grant to D.L.G., and an Alliance collaborative grant to D.L.G, M.M.H and S.M and graduate student funding awarded to J.A.B. Thank you to those who contributed to our crowdfunding campaign, who helped us fund preliminary pilot rodent studies. Thank you to April Mahovlic for her assistance characterizing the *Muc2*^+/-^ and *Muc2*^-/-^ models. Thank you to Christian Mustapich for his assistance with light/dark video analysis. Thank you to Ryan Bonnie and Helen Chiang for their assistance during mouse experiments. Thank you to AC for their intellectual contribution early in the project. Thank you to Innovate Phytoceuticals (Kelowna, BC, Canada) for their analysis of tryptophan and indole metabolites in serum. Thank you to Dr. Wesley Zandberg for the use of his lyophilizer for cecal samples used in SCFA analysis. Figure 5 was created in https://BioRender.com.

## Author contributions

D.L.G funded the experiment. M.M.H and D.L.G conceived the initial research question. J.A.B and D.L.G designed the experiments. J.A.B collected and analyzed all data. N.H calculated the Average American Diet dose used throughout the experiment. J.K.J assisted with behavioral and metabolic mouse experiments, data analysis and scored colon tissues. R.I. and E.Y assisted with paper revisions including ZO-2 staining. J.G. contributed experimental protocols used for open field behavior testing. M.L.B blindly scored open field experiments. C.J.M assisted with qPCR. A.V. and J.Y. assisted with mouse experiments, genotyping and metabolic experiments. A.V. also served as a scorer for colon tissue samples and cut and stained tissues for analysis. M.M.J and K.K.S assisted with corticosterone analysis and provided input regarding experimental design. S.G. provided protocols for glucose and insulin tolerance testing and assisted with metabolic analysis interpretation and allowed use of his GC for SCFA analysis. C.L and A.C pulled peaks and performed GC analysis for SCFA analysis. S.J.M and R.T.G. created the method used to detect glyphosate. D.L.G and J.A.B wrote the manuscript. All authors edited and approved the final manuscript.

