## Supplemental Materials for "Prenatal exposure to dietary levels of glyphosate disrupts metabolic, immune, and behavioral markers across generations in mice"

table S1: Changes made to original pre-registration plan and rationale

| Design Plan | Original 2020 Pre-registration | Changes Made | Reason for Change |
| --- | --- | --- | --- |
| Blinding | No blinding is involved in this study. | Experiments were blinded during all analysis. Samples were randomly assigned a number (1-169) and were not decoded until analysis had been completed. | Reduce risk of bias. |
| Randomization | No response | Samples were randomly assigned using a random number generator. | Reduce risk of bias. |
| Data collection procedures | For insulin tolerance testing, animals will not have fasted to prevent hypoglycemia. | Animals were fasted. | A dose that did not induce hypoglycemia in our *Muc2*-/- animal model was found by another researcher in our lab. Therefore, animals were fasted to allow for more accurate comparisons between animals and groups. |
| Data collection procedures | If at any time an animal goes hypoglycemic (blood glucose reading below 3mmol/L), the animal will receive glucose | Replaced glucose with sucrose and replaced gavage with IP injection. | UBC Animal Care suggested replacing glucose with sucrose and gavage with IP injection as it gets into the system faster and can recover animals more effectively. |
| Data collection procedures | 8 male *Muc2*^-/+^ and 8 female *Muc2*^-/+^ mice will be randomly selected using a random number generator and mated. 4 breeding pairs will be exposed to EPA dose glyphosate (1.75mg/kg body weight/day) while the other 4 breeding pairs will receive autoclaved reverse osmosis water. | Changed breeding scheme to *Muc2*^-/+^ and *Muc2*^-/-^ and increased breeding pairs to meet target n numbers. An additional 0.01mg/kg/day dose was added. | Difficulties reaching target n number in the original pilot study. Additionally, some effects observed using the low dose in direct exposure suggested this dose may elicit effects, so we added it to generational studies. |
| Data collection procedures | Animals will be subjected to the open field test. | Addition of light/dark, novel object and radial arm maze test | Access to new mazes. |
| Data collection procedures | The videotape will be analyzed using software developed by Dr. Issac Li to reduce experimental bias. | Purchased EthoVision software for data analysis. | Access to commercially available, validated software. |
| Sample Size | Previous microbiome studies conducted in the *Muc2*^-/-^ animal model have shown that an n = 12 (6 male and 6 female) per group provides adequate statistical power. | The sample size is highly variable across genotypes, sex, and group, with many groups failing to meet the n=12 | Variations/difficulties with breeding. |
| Data collection procedures | Animals will be subjected to the open field test. | Addition of light/dark, novel object and radial arm maze test | Access to new mazes. |
| Data collection procedures | The videotape will be analyzed using software developed by Dr. Issac Li to reduce experimental bias. | Purchased EthoVision software for data analysis. | Access to commercially available, validated software. |
| Sample Size | Previous microbiome studies conducted in the *Muc2*^-/-^ animal model have shown that an n = 12 (6 male and 6 female) per group provides adequate statistical power. | The sample size is highly variable across genotypes, sex, and group, with many groups failing to meet the n=12 | Variations/difficulties with breeding. |
| Sample size rationale | No response | Mead's Resource Equation | A power analysis could not be conducted due to a lack of similar studies in the field. However, discovered Mead's Resource Equation after pre-registration which was used to obtain sample size estimate. |

table S1: Changes made to original pre-registration plan and rationale

| Design Plan | Original 2020 Pre-registration | Changes Made | Reason for Change |
| --- | --- | --- | --- |
| Manipulated variables | Genotype (*Muc2*^-/-^, *Muc2*^-/+^ and or *Muc2*^+/+^) | It was discovered that *Muc2*-/+ and *Muc2*+/+ are the same regarding colitis resistance, metabolic effects, and behaviour, so we decided to alter the breeding scheme. | Maximize n number. |
| Data Exclusion | Outliers will be included in the analysis. | Outliers were removed from the study. | Some data sets featured extreme outliers far outside of normal biological variability. These outliers resulted in an extreme skew of the data. |
| Exploratory analysis | No response | Specific findings, including decreased locomotor activity, led to further exploratory analysis, including alpha synuclein quantification. | These were not expected findings and are reported as being exploratory. Additionally, increased work conducted in the field over the past four years led to increased knowledge surrounding the potential link between PD and glyphosate. Any future work looking at the link between glyphosate and locomotor activity (e.g., Parkinson's disease) will be registered under a new pre-registration. |
| Measured variables | Histology will be scored by two blind individuals concerning animal treatment and genotype. | Three individuals scored Histology. | Reduce bias/subjective scoring. |
| Statistical Models | Comparisons between means (i.e., cytokine data) will be analyzed using a one-way ANOVA. | Comparisons between means were analyzed using different appropriate methods. | Improved knowledge led to more stringent and robust methods being used. Additionally, a small sample size obtained in many groups and a large amount of variability within the data resulted in alterations to statistical methods. |
| Indices | Cytokine data will be analyzed using several stable reference genes including Beta Actin, 18s, and *Tbp*, and data will be normalized to the healthy wild-type control group. | Cytokine data was normalized to reference genes *Eef2* and T*bp*. Additionally, samples were not normalized to the control group. | 18s and Beta Actin are unstable within the gut during inflammation. The decision was made to use more stable reference genes. |
| Transformations | No response | If data exhibited log-normal distribution, it was log-transformed. | Improved knowledge led to more stringent and robust methods being used. Additionally, a small sample size obtained in many groups and a large amount of variability within the data resulted in alterations to statistical methods. |
| Inference Criteria | We will use the standard p<.05 criteria to determine if the ANOVA and the post hoc test suggest that the results significantly differ from those expected. | Comparisons between means were analyzed using different appropriate methods. | Improved knowledge led to more stringent and robust methods being used. Additionally, a small sample size obtained in many groups and a large amount of variability within the data resulted in alterations to statistical methods. |

table S2: Breakdown of sex and genotype combinations within parental generation mice.

| Group | Number of *Muc2*^+/-^ Males | Number of *Muc2*^+/-^ Females | Number of *Muc2*^-/-^ Males | Number of  *Muc2*^-/-^ Females |
| --- | --- | --- | --- | --- |
| No Exposure | 3 | 3 | 3 | 3 |
| Average American Diet | 2 | 4 | 4 | 2 |
| EPA Upper Limit | 4 | 2 | 2 | 4 |

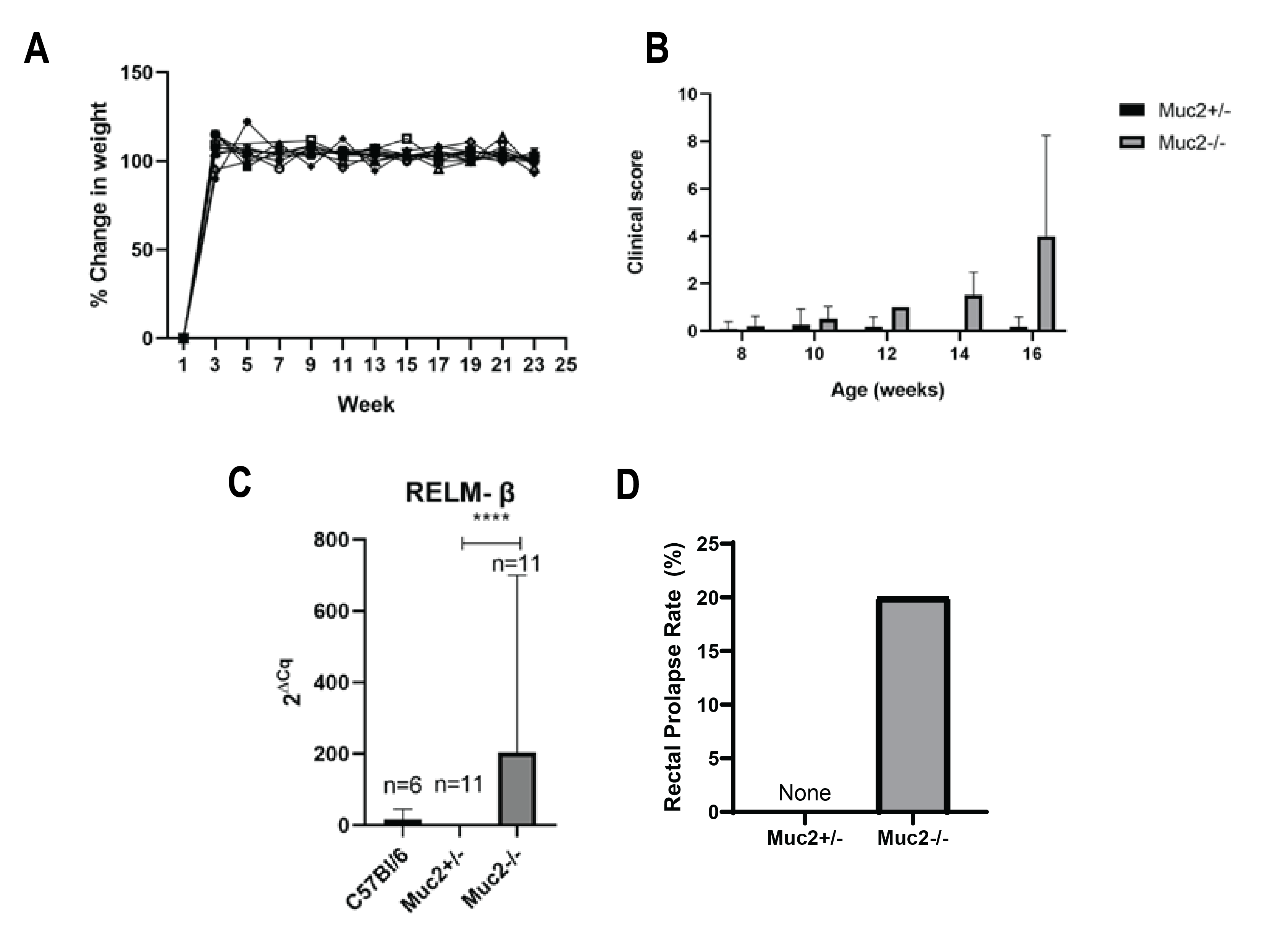

**figure S1:** **[A]** Scatter plot showing % change in weight versus week of study for 11 *Muc2*^+/-^ mice. Mice were each weighed every 2 weeks. During the first week of study mice were between 5 and 8 weeks of age, and at the 23^rd^ week of study, mice were between 27 and 30 weeks of age. **[B]** Clinical scores were recorded every 2 weeks for *Muc2*^+/-^ mice (n=11), and every week for *Muc2*^-/-^ mice (n=10). Scores for *Muc2*^-/-^ mice were averaged over 2 weeks to be consistent with *Muc2*^+/-^ measurements. Data for *Muc2*^+/-^ mice is contained in black boxes, and data for *Muc2*^-/-^ mice is in gray boxes. **[C]** Bar plot demonstrating difference in Cq values of RELMβ for C57Bl/6, *Muc2*^+/-^ and *Muc2*^-/-^ animals. ∆Cq was calculated as the difference between the reference gene and the target gene. **[D]** Bar plot showing occurrences of rectal prolapse during study of *Muc2*^+/-^ mice and *Muc2*^-/-^ mice. Rectal prolapse occurs in 40% of *Muc2*^-/-^ mice by 6 months of age. *Muc2*^+/-^ mice (n=11) were monitored over a period of 6 months, whereas *Muc2*^-/-^ mice (n=10) were monitored over a period of 4 months.

table S3: Colonic histological scoring criteria used for healthy and colitis-susceptible litter mates.

| *Muc2*^+/-^, *Muc2*^+/+^ and *Muc2*^-/-^ Combined Scoring | | | |
| --- | --- | --- | --- |
| Category | **Criterion** | **Definition** | **Score** |
| Inflammatory Cell Infiltrate | Severity and Extent | ***Density and expansion of leukocytes*** | |
|  |  | **Minimal:** Submucosa | 1 |
|  |  | **Mild:** Submucosa sometimes extending into mucosa | 2 |
|  |  | **Moderate:** Submucosa and mucosal | 3 |
|  |  | **Marked:** mucosal, submucosal, and transmural | 4 |
| Hyperplasia | Elongated Crypts | **Minimal:** <25% Compared to Healthy Control | 1 |
|  |  | **Mild:** 25-35% | 2 |
|  |  | **Moderate:** 36-50%, mitoses in upper third of crypt epithelium, distant from crypt base | 3 |
|  |  | **Marked:** >51%, mitoses in upper third of crypt epithelium, distant from crypt base | 4 |
| Epithelial Changes | Cryptitis | Neutrophils between crypt epithelial cells | 1 |
|  | Cryptitis, crypt ulceration | Neutrophils in crypt lumen | 2 |
|  | Focal erosions | Loss of surface epithelium | 3 |
|  | Crypt Destruction | Loss of crypts | 4 |
| Blood |  | Absent | 0 |
|  |  | Present | 2 |
| Crypt Abscess (severe) |  | 1 for each | 1/each |
| Edema |  | Absent | 0 |
|  |  | Moderate | 1 |
|  |  | Marked | 2 |

table S4: Additional histology scoring criteria used for healthy animals

| *Muc2*^+/-^, *Muc2*^+/+^ Only - Goblet Cell Depletion | | | |
| --- | --- | --- | --- |
| Category | **Criterion** | **Definition** | **Score** |
| Goblet Cell Depletion |  | ***Reduction of goblet cell numbers relative to baseline goblet cell numbers per crypt*** | |
|  |  | **Minimal:** <20% | 1 |
|  |  | **Mild:** 21-35% | 2 |
|  |  | **Moderate:** 36-50% | 3 |
|  |  | **Marked:** >50% | 4 |

The scoring scheme was adapted from the studies by Haskey et al., (2022)^71^ and Erben et al. (2014).^72^

table S5: Primer sequences used

| Name | Gene | Forward | Reverse | Efficiency |
| --- | --- | --- | --- | --- |
| aSyn | *Snca* | gcaagggtgaggaggggta | cctctgaaggcatttcataagcc | 102% |
| CAT3 | *Cat3* | ggacgctcagcttttcattc | ttgtccagaagagcctggat | 101% |
| Chymase 1 | *Cma1* | ataagcctaaggcccaaatatga | caatgatctctccagctttggt | 94% |
| EEF2 | *Eef2* | tgtcagtcatcgcccatgtg | catccttgcgagtgtcagtga | 110% |
| FOXp3 | *Foxp3* | cctctgccgttatccagcctgcc | gcccttgggtgcagtcttcc | 101% |
| GPX3 | *Gpx3* | gatgtgaacggggagaaaga | ttcatgggttcccaaaagag | 101% |
| IDO1 | *Ido1* | caaagcaatccccactgtatcc | acaaagtcacgcatcctcttaaa | 103% |
| IFN-ϒ | *Ifng* | tcaagtggcatagatgtggaagaa | tggctctgcaggattttcatg | 109% |
| IL-10 | *Il10* | agggccctttgctatggtgt | tggccacagttttcagggat | 90% |
| IL-17a | *Il17a* | tccgaggagtcagtgctaaa | tccgaggagtcagtgctaaa | 100% |
| IL-1ϐ | *Il1b* | gccaccttttgacagtgatgag | caaaggtttggaagcagcccttc | 109% |
| IL-22 | *Il22* | agctcctgtcacatcagcg | agcttcttctcgctcagacg | 103% |
| iNOS | *Nos2* | gacattacgacccctcccac | gcacatgcaaggaagggaac | 94% |
| MCP1 | *Ccl2* | gcagcaggtgtcccaaagaa | atttacgggtcaacttcacattcaa | 100% |
| MCPT1 | *Mcpt1* | ttccaggtctgtgtgggaag | tccagggcacatatgcagag | 100% |
| MCPT2 | *Mcpt2* | aacggttcagaaggagaggtg | tctgtgtgtgggttcgttc | 108% |
| *MUC2* | *Muc2* | gccagatcccgaaacca | tataggagtctcggcagtca | 97% |
| Reg3ϒ | *Reg3g* | cccgtataaccatcaccatcat | ggcatctttcttggcaacttc | 100% |
| RELMβ | *Retnlb* | atgggtgtcactggatgtgctt | agcactggcagtggcaagta | 110% |
| TBP | *Tbp* | accgtgaatcttggctgtaaac | gcagcaaatcgcttgggatta | 110% |
| TGFβ | *Tgfb1* | gaccgcaacaacgccatcta | agccctgattccgtctcctt | 100% |
| TNFα | *Tnf* | catcttctcaaaattcgagtgacaa | tgggagtagaacaaggtacaaccc | 90% |
| SNAP25 | *Snap25* | caactggaacgcattgaggaa | ggccactactccatcctgattat | 90% |
| GFAP | *Gfap* | ccctggctcgtgtggattt | gaccgataccactcctctgtc | 96% |

**Glyphosate measurement in mouse chow and commercially available food items**

We developed an ultra-performance liquid chromatography (UPLC) method for measuring glyphosate in food. Briefly, dry samples were ground in a blender and 2.5 g of sample was weighed and spiked with isotopically labeled internal standard (Glyphosate-2-^13^C,^15^N, Sigma) followed by 5 mL of 1% formic acid in water. Samples were sonicated for 1 hour and then placed into a shaker incubator overnight at 200 RPM at 30^o^C. The next day, samples were centrifuged at 4000 RPM for 10 minutes and supernatant collected. The supernatant was filtered at 0.22 μm and an aliquot of the filtrate was diluted 1:2 with acetonitrile (ACN) and shaken. This was then transferred into an amber autosampler vial. Glyphosate was separated and identified with a Acquity I-Class UPLC using a Torus DEA 1.7μm 2.1 x 100 mm column (Waters, Mississauga). Mobile phase A consisted of 50mM ammonium formate at pH 2.9 (adjusted with formic acid) and mobile phase B consisted of acetonitrile + 0.9% formic acid. Stock standard solutions were prepared by dissolving 500mg glyphosate in 10mL 50:50 ACN:water in a 15mL centrifuge tubes sonicated for 20 minutes. Standards were sterilized by filtration through a 0.22 µm syringe filter. A standard curve was generated by performing an initial 1:50 dilution with 50:50 ACN:water as the diluent. Each successive dilution used a 1:2 dilution where 250 µL of the previous standard was mixed with 230 µL 50:50 ACN:water and 20 µL 5 mM sodium EDTA. 100µL of each standard was then transferred into autosampler vials with conical inserts. For both samples and standards, 10µL was injected. Six-point standard curves were run from 2.95 ug/mL to 0.092 ug/mL (figure S3, A).

Compounds were measured by tandem mass spectrometry (MS/MS) (Xevo TQ-S mass spectrometer in ESI negative mode, Waters). For MS/MS detection, the ion source had the following conditions: cone voltage of 30 V, capillary voltage 2.5 kV, desolvation temperature 600°C, desolvation gas (N_2_) flow 1000L/hr, cone gas (N_2_) flow 300 L/hr. In all cases ultra-high purity argon at 7 psi was used for the collision gas. For glyphosate, multiple reaction monitoring (MRM) was utilized. In brief, the precursor ion for glyphosate was 168 m/z, and the product ions of 62.9 m/z (quantifier) and 80.9 (qualifier) were monitored with collision energies of 16 and 18 eV, respectively. For the isotopically labeled internal standard, transitions of 170 > 62.9 and 170 > 80.9 were monitored.

Selectivity was indicated by complete baseline chromatographic separation of glyphosate from coeluting species (figure S3, B). Manual inspection of each chromatogram found no coeluting species in MRM mode. Linearity was determined from the six-point standard curve, in all cases linearity was 0.998 or greater (figure S3, A). The U.S. Food and Drug Administration (FDA, 1998) definitions for limit of detection (LOD) and lower limit of quantification (LLOQ) were explored. Method LOD was calculated as 0.08 μg/mL and method LLOQ was 0.34 μg/mL in extract solution. Interday precision (percent relative standard deviation (%RSD)) for glyphosate spiked method development samples (potato starch spiked with 0.10 μg/g glyphosate) was 7.21 % RSD. At this concentration, spike recoveries of 95.67 ± 6.04 % were achieved.

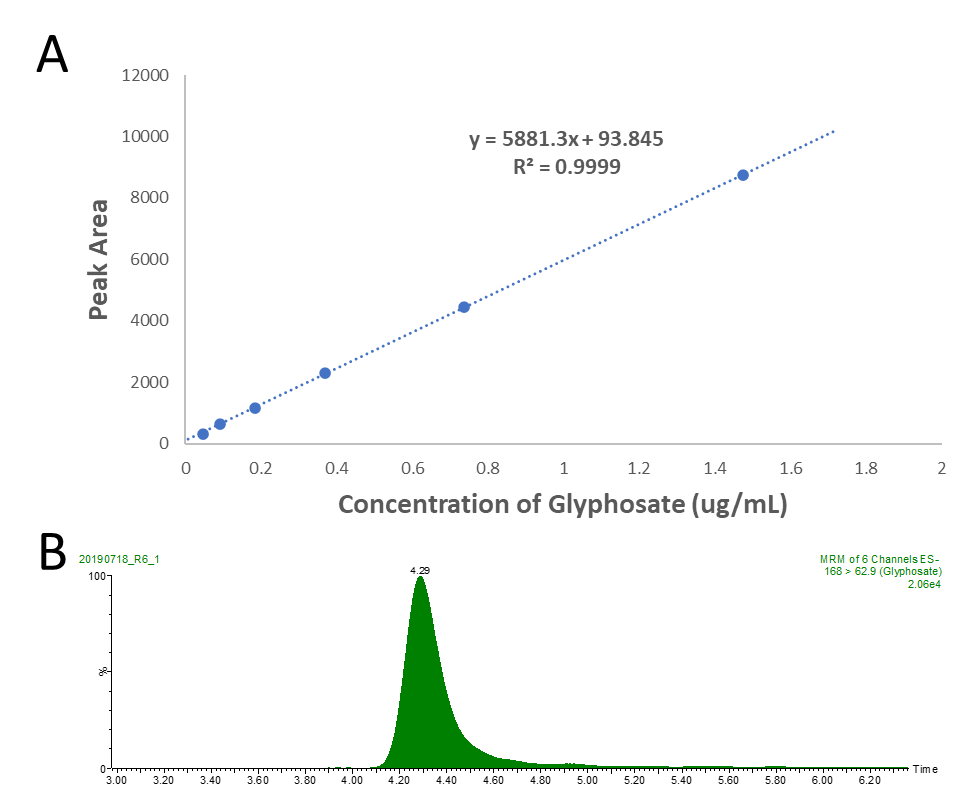

**figure S2:** A six-point standard curve of glyphosate demonstrating linearity, and **B** chromatogram of glyphosate detected in a sample demonstrating satisfactory selectivity and peak shape.

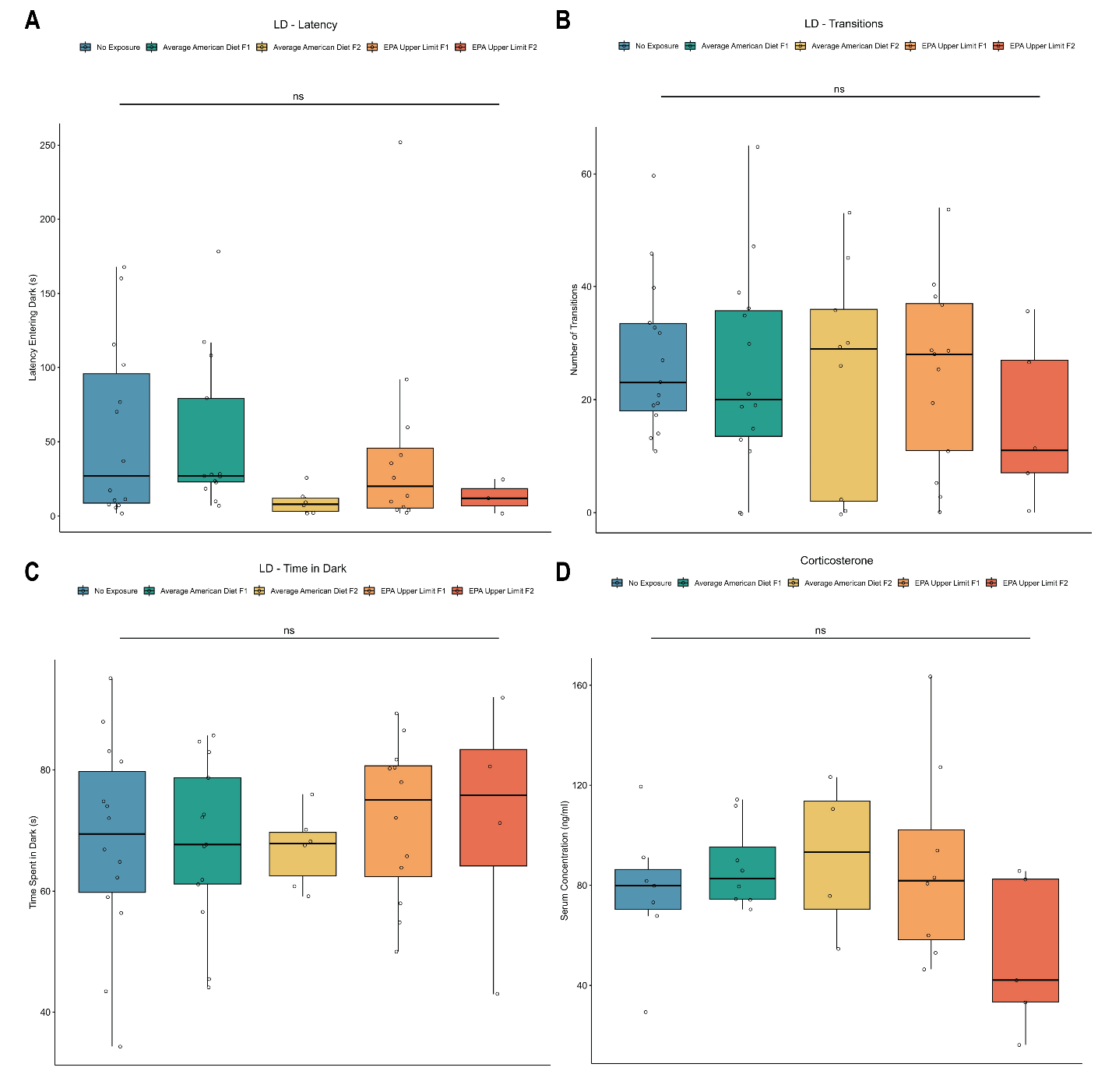

figure S3: Colitis-susceptible descendants of mice exposed to glyphosate do not exhibit signs of anxiety-like behavior during light/dark trial.

A) Latency was defined as the amount of time taken before a mouse entered the dark compartment. No significant differences were observed in latency entering the dark (Kruskal-Wallis, p=0.00783, τ_b_=-0.0835). Sex was not found to be a significant factor in latency entering dark (PERMANOVA, p=0.869). B) The number of transitions between the light and dark compartments were measured. No significant differences were observed in the number of transitions between the two compartments (ANOVA, p=0.682, ω^2^=0). Sex was not found to be a significant factor in transitions between the light and dark compartments (PERMANOVA, p=0.826). C) No significant differences were observed regarding the amount of time a mouse spent within the dark compartment (Kruskal-Wallis, p=0.336, τ_b_=-0.147). Sex was not found to be a significant factor for time spent within each compartment (PERMANOVA, p=0.616). Each treatment group was compared to its no exposure control group shown in blue. Each point represents an individual mouse. The bottom and top of the boxes are the first and third quartiles, the middle band inside the boxes is the median and the whiskers contain the upper and lower 1.5 interquartile range (IQR). Serum corticosterone levels were measured using ELISA. No significant differences were observed between any of the treatment groups and control (ANOVA, p=0.569, ω^2^ = 0). Sex was not found to be a significant factor (PERMANOVA, p=0.452). Each treatment group was compared to its no exposure control group shown in blue. Each point represents an individual mouse. The bottom and top of the boxes are the first and third quartiles, the middle band inside the boxes is the median and the whiskers contain the upper and lower 1.5 interquartile range (IQR).

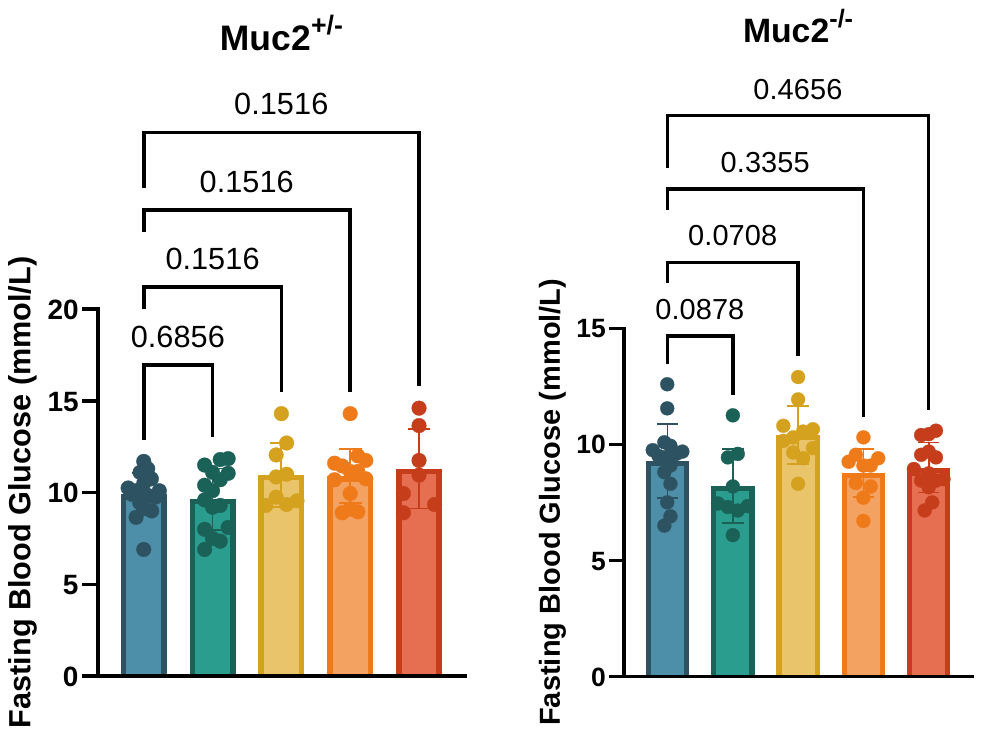

figure S4: Healthy (Muc2^+/-^) and colitis-susceptible (Muc2^-/-^) mice exhibit no significant differences in fasting glucose levels prior to OGTT or ITT (Muc2^+/-^, ANOVA, *p*=0.0541, FDR-correction; Muc2^-/-^, ANOVA, *p*=0.0078).

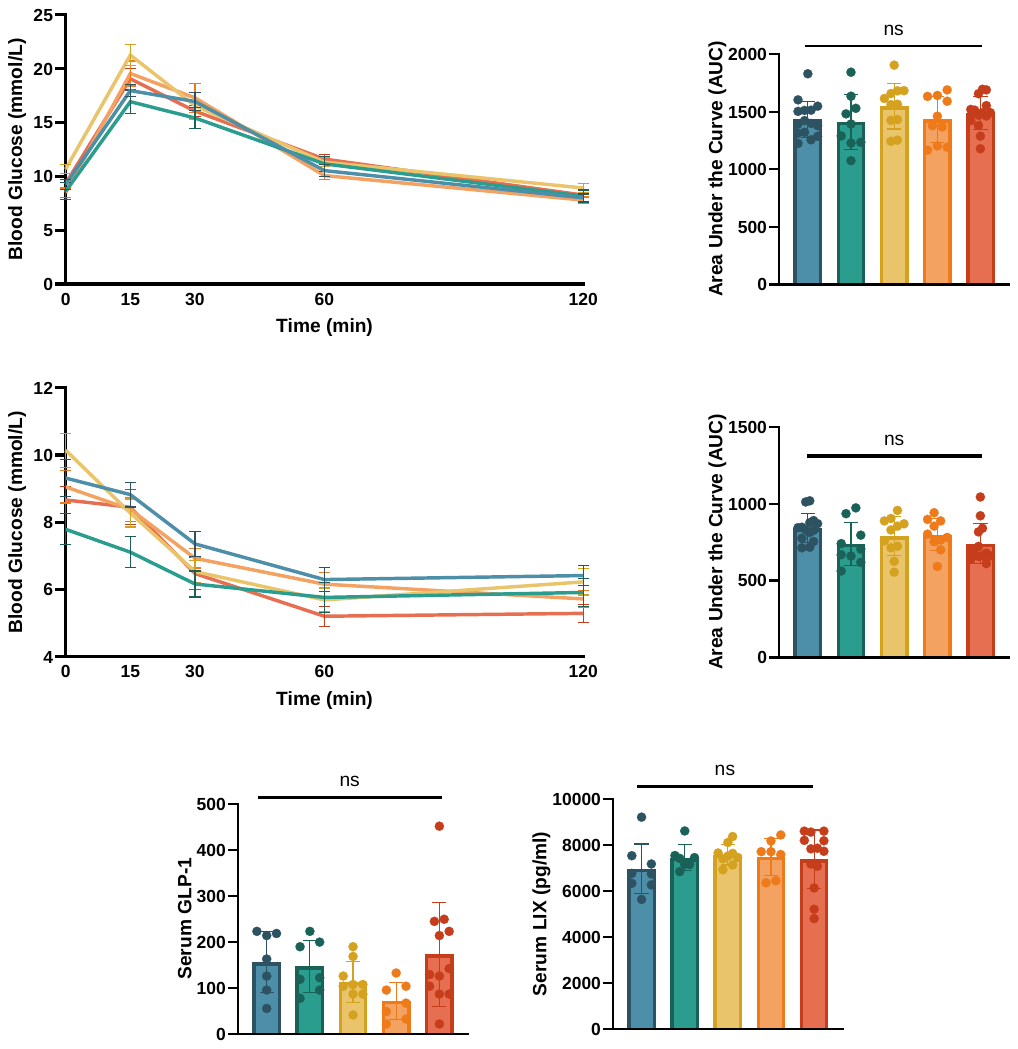

figure S5: Transgenerational exposure to glyphosate does not alter glucose metabolism in colitis-susceptible offspring. [A]

Animals were fasted for six hours before receiving a 2g/kg fasted body weight oral gavage of sterile filtered glucose. Blood glucose measurements were taken before and 15-, 30-, 60- and 120 minutes following gavage. Each treatment group was compared to its no exposure control group shown in blue. No significant differences were observed between any treatment group at any point in time (Two-way ANOVA). Each point represents the average blood glucose reading for the group at a given time point. No significant differences were observed between treatment groups for the area under the curve (ANOVA, p=0.475, ω^2^ = 0). Sex was not found to be an influencing factor (PERMANOVA, p=0.533). Each point represents an individual mouse. [B] Ancestral glyphosate exposure does not significantly influence insulin tolerance in colitis-susceptible descendants.

animals were fasted for six hours before receiving a 0.5U/kg fasted body weight intraperitoneal injection of Insulin. Blood glucose measurements were taken before and 15-, 30-, 60- and 120 minutes following injection. Each treatment group was compared to its no exposure control group shown in blue. No significant differences were observed between any treatment group at any point in time (Two-way ANOVA). Each point represents the average blood glucose reading for the group at a given time point. No significant differences were observed between treatment groups for the area under the curve (ANOVA, p=0.117, ω^2^ = 0.06). Sex was not found to be a contributing factor (PERMANOVA, p=0.095). Each point represents an individual mouse.

table S6: Full test statistics, confidence intervals, and sample sizes broken down by sex.

| Figure Panel | 95% CI (Range) | Sample Size n(sex) | Figure Panel | 95% CI (Range) | Sample Size n(sex) |
| --- | --- | --- | --- | --- | --- |
| 1A | 2.832 to 3.856; 4.889 to 6.236; 4.291 to 5.375; 4.333 to 5.513; 5.153 to 10.35 | 16 [10F, 6M], 16[6F, 10M], 6[3F, 3M], 13, [6F, 7M], 5[2F, 3M] | **3A (Distance)** | 3,955 to 4,825; 4,164 to 4,879; 3,820 to 4,597; 3,287 to 4,265; 2,127 to 3,406 | 14[9F, 5M], 15[6F, 9M), 9[7F, 2M], 9[6F, 3M] 12[4F, 8M] |
| 1B (Goblet Cell) | 1.590 to 2.285; 0.9296 to 1.820; 0.5153 to 1.235; 1.671 to 2.679 | 16 [10F, 6M], 16[6F, 10M], 6[3F, 3M], 13[6F, 7M], 5[2F, 3M] | **3A (Velocity)** | 6.604 to 8.059; 6.946 to 8.155; 6.376 to 7.691; 5.488 to 7.130; 4.201 to 6.524 | 14[9F, 5M], 15[6F, 9M), 9[7F, 2M], 9[6F, 3M] 12[4F, 8M] |
| 1B (Hyperplasia) | 0.4025 to 1.097; -0.1240 to 0.7668; 0.6046 to 1.324; 1.046 to 2.054 | 16 [10F, 6M], 16[6F, 10M], 6[3F, 3M], 13[6F, 7M], 5[2F, 3M] | **3B (Total Rearing)** | 89.03 to 112.5; 97.56 to 115; 91.14 to 104.2; 84.88 to 100.9; 52.33 to 72.5 | 14[9F, 5M], 15[6F, 9M), 9[7F, 2M], 9[6F, 3M] 12[4F, 8M] |
| 1C (Goblet Cell) | 0.06603 to 0.3625; 0.6687 to 1.769; 1.500 to 1.500; 1.008 to 2.278; 0.9341 to 4.266 | 16 [10F, 6M], 16[6F, 10M], 6[3F, 3M], 13[6F, 7M], 5[2F, 3M] |  |  |  |
| 1C (Mucin2) | 0.05558 to 0.2313; 0.01102 to 0.08596; 0.00 to 0.05721; 0.008579 to 0.07219; 0.007208 to 0.02817 | 12[6F, 6M], 9[4F, 5M], 7[4F, 3M], 10[5F, 5M], 7[3F, 4M] | **3B (Unsupported Rearing)** | 34.02 to 51.40; 38.89 to 49.51; 34.07 to 48.60; 32.81 to 52.30; -2.965 to 16.97 | 14[9F, 5M], 15[6F, 9M), 9[7F, 2M], 9[6F, 3M] 12[4F, 8M] |
| 1D | 12.31 to 23.78; 16.42 to 24.05; 9.549 to 55.68; 17.01 to 49.12; 18.20 to 76.73 | 13[7F, 6M], 18[7F, 11M], 4[2F, 2M], 12[6F, 6M], 5[2F, 3M] |  |  |  |
| 1F | 23.8 to 57.96; 10.15 to 48.78; 36.89 to 80.85; 23.30 to 102; 65.21 to 143 | 13[7F, 6M], 7 [3F, 4M], 12[10F, 2M], 9[3F, 6M], 13[6F, 7M] | **3C** | 5.880 to -1.587; -2.316 to 2.461; -3.228 to -0.6248; -2.255 to 2.038 | 14[9F, 5M], 15[6F, 9M], 9[7F, 2M], 9[6F, 3M], 12[4F, 8M] |
| 1H | -1.616 to -1.018; -1.181 to -0.06023; -0.7883 to 0.3852; -1.690 to -0.2432; -1.059 to -0.04305 | 16[8F, 8M], 10[4F, 6M], 10[6F, 4M], 10[4F, 6M], 13[6F, 7M] | **3D** | 0.01341 to 0.01892; 0.007864 to 0.01485; 0.01207 to 0.01785; 0.01155 to 0.02302; 0.005679 to 0.01532 | 11 [8F, 3M], 7[3F, 4M], 5[2F, 3M], 7[3F, 4M], 5[3F, 2M] |
| 1I | -0.9501 to -0.2288; -0.6798 to -0.03084; -0.1371 to 0.7111; -0.7527 to 0.2975; 0.02421 to 0.5396 | 16[8F, 8M], 10[4F, 6M], 10[6F, 4M], 10[4F, 6M], 12[6F, 6M] | **3E (ASYN)** | -0.6582 to -0.2436; -0.6516 to -0.06955; -0.3153 to 0.8006; -0.8155 to 0.3696; -0.3821 to 0.4631 | 16[8F, 8M], 10[4F, 6M], 10[6F, 4M], 10[4F, 6M], 13[6F, 7M] |
| 1J | 0.04258 to 0.1064; 0.02207 to 0.3082; 0.2886 to 1.111; 0.002768 to 0.1858; 0.2096 to 0.8429 | 14[7F, 7M], 9[4F, 5M], 10 [6F, 4M], 8[4F, 4M], 13[6F, 7M] |  |  |  |
| 1K | 0.4108 to 0.9766; 0.4072 to 1.114; 3.283 to 28.71; 0.2849 to 1.379; 1.216 to 9.054 | 16[8F, 8M], 8[4F, 4M], 10[6F, 4M], 8[4F, 4M], 12[6F, 6M] | **3E (GFAP)** | 0.3332 to 0.5350; 0.2088 to 0.9642; 0.4237 to 3.697; 0.3162 to 0.9536; 0.6091 to 5.532 | 16[8F,8M], 8[3F, 5M], 12[5F, 7M], 10[4F, 6M], 10[5F, 5M] |
| 2A | 1,386 to 1,600; 1,362 to 1,541; 1,383 to 1,609; 1,523 to 1,709; 1,490 to 2,197 | 16[10F, 6M], 16[6F, 10M], 9[5F, 4M], 13[6F, 7M], 7[3F, 4M] | **3E (SNAP-25)** | -0.5740 to -0.2797; -0.4467 to -0.008742; -0.3578 to 0.5369; -0.6208 to -0.2852; -0.4385 to 0.8979) | 12[6F,6M], 5[2F, 3M], 8[4F, 4M], 3[2F, 1M], 11[6F, 5M] |
| 2B | 804.9 to 942.1; 832 to 942.7; 775.6 to 1,012; 909.1 to 1,120; 706.4 to 952.6 | 16[10F, 6M], 16[6F, 10M], 9[5F, 4M], 13[6F, 7M], 7[3F, 4M] | **4B** | 128.5 to 179.6; 180.4 to 242.7; 102.8 to 255.7; 147.2 to 242.6; 144.3 to 256.4 | 16 [10F, 6M], 16[6F, 10M], 6[3F, 3M], 9[5F, 4M], 8[4F, 4M] |
| 2C | 31.11 to 193.1; -29.28 to 101.6; 16.77 to 161.6; 105 to 232.7 | 8[3F, 5M], 8[4F, 4M], 5[2F, 3M], 11[5F, 6M], 6[3F, 3M] | **4D (Aerobic)** | 0.07717 to 0.1471; 0.07931 to 0.1763; 0.07840 to 0.1651; 0.1500 to 0.1898; 0.1212 to 0.1883 | 14 [8F, 6M], 7[3F, 4M], 10[5F, 5M], 10[4F, 6M], 11[5F, 6M] |
| 2D | 3.866 to 10.21; 2.068 to 3.745; 2.432 to 6.528; 1.905 to 3.622; 2.408 to 5.287 | 8[3F, 5M], 9[4F, 5M], 5[2F, 3M], 11[5F, 6M], 8[4F, 4M] | **4D (Pathogenic)** | 0.3614 to 0.5330; 0.4263 to 0.6698; 0.4216 to 0.5744; 0.5655 to 0.6271; 0.5216 to 0.6139 | 14 [8F, 6M], 7[3F, 4M], 10[5F, 5M], 10[4F, 6M], 10[5F, 5M] |
| 2E | 144.2 to 738.2; 554.1 to 1,863; 441.5 to 1,605; 670.6 to 2,796; 280.1 to 853.1 | 8[3F, 5M], 9[4F, 5M], 5[2F, 3M], 11[5F, 6M], 8[4F, 4M] | **4D (Biofilm)** | 0.09267 to 0.1564; 0.1110 to 0.2023; 0.09516 to 0.1844; 0.1652 to 0.2152; 0.1353 to 0.2006 | 14 [8F, 6M], 7[3F, 4M], 10[5F, 5M], 10[4F, 6M], 11[5F, 6M] |
| 2F | 5,975 to 6,917; 7,861 to 8,184; 6,447 to 8,646; 6,864 to 8,529; 6,556 to 7,813 | 8[3F, 5M], 6[3F, 3M], 5[2F, 3M], 11[5F, 6M], 8[4F, 4M] |  |  |  |
